# Electrostatic Complementarity at the ClpX Substrate-Entry Channel Governs ATP-Driven Protein Unfolding

**DOI:** 10.64898/2026.07.31.739446

**Authors:** Yifei Lyu, Naseer Iqbal, Alireza Ghanbarpour

## Abstract

AAA+ proteases maintain proteostasis by mechanically unfolding protein substrates before degradation, yet how ATP-driven pulling is converted into productive unfolding remains poorly understood. Here, using ssrA-tagged GFP substrates spanning a range of surface charges, we show that electrostatic complementarity between the positively charged ClpX substrate-entry channel and folded substrate domains governs unfolding efficiency. Positively charged substrates unfolded inefficiently despite substrate recognition, thermal stability, ATPase activation, and pore-loop engagement comparable to negatively charged substrates. Cryo-EM structures of a positively charged substrate revealed multiple substrate-engagement states arising from electrostatic incompatibility at the ClpX substrate-entry channel, whereas negatively charged substrates formed favorable electrostatic interactions that likely stabilize partially unfolded intermediates during successive ATP-driven unfolding attempts. Together, these findings identify electrostatic complementarity as a key determinant of productive ATP-driven protein unfolding and suggest that tuning substrate or substrate-entry electrostatics provides a mechanism for regulating AAA+ protease activity.

## INTRODUCTION

AAA+ proteases are ATP-powered molecular machines that carry out much of intracellular protein degradation in bacteria and human mitochondria ^1–2^. These enzymes harness the energy of ATP hydrolysis to mechanically unfold protein substrates and translocate the resulting polypeptides into an enclosed proteolytic chamber for degradation^3^. Because stable protein domains must be mechanically unfolded before degradation, the efficiency of substrate unfolding is a major determinant of AAA+ protease activity. Failure of these pathways results in the accumulation of toxic protein aggregates, impaired cellular viability, and numerous diseases associated with proteostasis collapse^2, 4–5^.

The bacterial ClpXP protease is a paradigmatic AAA+ degradation machine that plays a central role in protein quality control and the degradation of regulatory proteins required for cellular adaptation ^3, 6^. Its mitochondrial homolog is essential for maintaining mitochondrial proteostasis and physiology, has been implicated in several human diseases^7^, and has recently emerged as a promising target for anticancer therapy^8–9^. In this system, the hexameric AAA+ unfoldase ClpX recognizes a short, typically unstructured peptide degradation signal (degron) and uses ATP hydrolysis to mechanically unfold the folded domain of protein substrates before translocating the resulting unfolded polypeptides into the tetradecameric ClpP peptidase for degradation (Fig. 1A)^3^. In bacteria, many ClpXP substrates are generated through the tmRNA quality-control pathway, which appends a short ssrA degron to incompletely translated proteins produced during ribosome stalling^10^. Because ssrA tagging can occur on a wide variety of proteins ^11^, ClpXP must efficiently process substrates with diverse structural and physicochemical properties.

**Figure 1.**
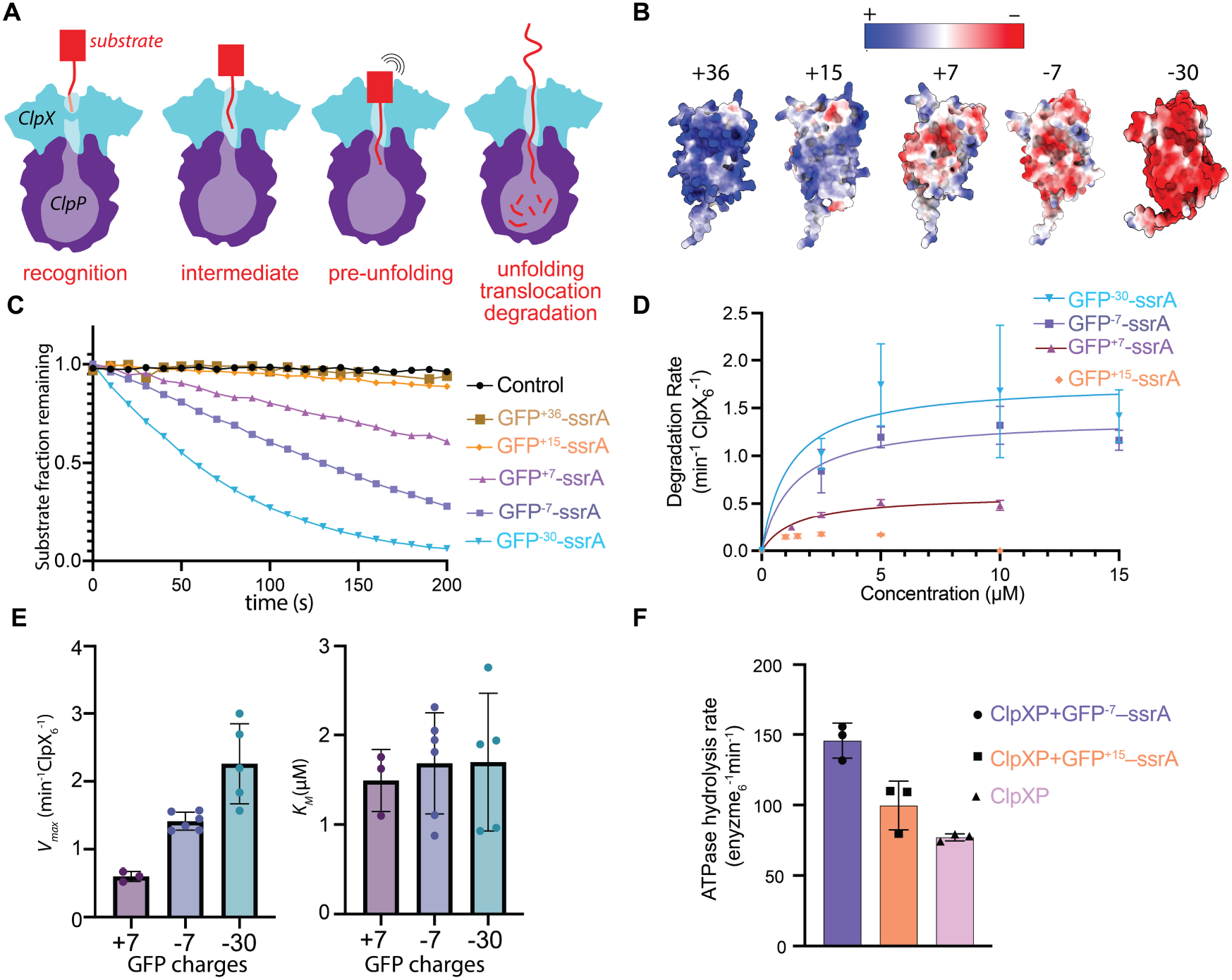
Substrate surface charge modulates ClpXP-mediated degradation. (**A**) Schematic model of ClpXP-mediated substrate processing. ClpX recognizes a protein substrate through its degron, engages the substrate in the axial channel, applies ATP-driven mechanical force to unfold the folded domain, and translocates the resulting polypeptide into ClpP for degradation. (**B**) Electrostatic surface representations of AlphaFold3-predicted GFP–ssrA variants with systematically varied net charge (+36, +15, +7, −7, and −30). Blue and red indicate positively and negatively charged surface regions, respectively. (**C**) Representative degradation time courses of GFP charge variants monitored by loss of GFP fluorescence during ClpXP-mediated degradation. Increasing substrate positivity progressively reduced degradation efficiency. Reactions contained 0.5 µM ClpX_6_, 1.5 µM ClpP_14_, and 5 µM GFP–ssrA substrate and were performed at 30 °C. GFP^+15^ incubated without enzyme served as the negative control. (**D**) Michaelis–Menten analysis of degradation kinetics for GFP charge variants. GFP^+15^–ssrA degradation was too slow to permit reliable determination of kinetic parameters and is therefore shown without fitting. (**E**) Derived kinetic parameters (*V*_*max*_ and *K*_*M*_) for GFP charge variants. Error bars represent standard deviations from independent experiments. (n = 3–6 independent experiments). (**F**) ATPase activity of ClpXP in the presence of GFP charge variants. Although GFP^+15^–ssrA was degraded inefficiently, it stimulated ATP hydrolysis above the basal activity of ClpXP.

Structural studies have established that ClpX adopts a shallow hexameric spiral in which the axial channel is lined by positively charged RKH (Arg^228^, Lys^229^, and His^230^), pore-1, and pore-2 loops that collectively mediate degron recognition, substrate engagement, unfolding, and translocation^12–16^. The RKH and pore-2 loops primarily contribute to degron recognition and initial substrate engagement^14, 17^, whereas the pore-1 loops provide the principal mechanical grip required for ATP-driven translocation and unfolding ^18–20^.

Recent cryo-EM structures of ClpXP stalled during substrate unfolding revealed that folded substrate domains are tightly encircled by positively charged RKH loops at the entrance of the ClpX axial channel^20^. Although these structures revealed extensive contacts between the folded substrate and the substrate-entry channel, whether these interactions actively promote ATP-driven unfolding or simply facilitate substrate positioning remained unknown. To address this question, we engineered a panel of supercharged GFP substrates bearing identical C-terminal ssrA degrons but spanning a broad range of surface charges and combined biochemical degradation assays, targeted RKH-loop mutagenesis, and cryo-EM to determine how substrate surface charge influences ClpXP-mediated unfolding. Together, these experiments demonstrate that electrostatic complementarity between the substrate surface and the positively charged ClpX substrate-entry channel is a key determinant of productive ATP-driven protein unfolding, revealing electrostatic complementarity as a previously unrecognized mechanism for regulating AAA+ protease activity.

## RESULTS

### Charge-Variant GFP Substrates Reveal Electrostatic Contributions to Productive Unfolding

Our previous structural analysis revealed that positively charged RKH loops encircle the folded domain of an engaged substrate at the entrance of the ClpX axial channel^20^ (Fig. S1). This observation raised the question of whether substrate electrostatics influence unfolding efficiency. To test this hypothesis, we appended a C-terminal ssrA degron to a panel of previously engineered supercharged GFP variants^21^ that differ systematically in net surface charge (+36, +15, +7, –7, or –30) through substitutions of solvent-exposed residues (Fig. 1B). We reasoned that substrates with opposing surface charges would exhibit different degradation efficiencies if electrostatic interactions contribute to productive unfolding. Degradation kinetics monitored by GFP fluorescence loss or SDS PAGE gel analysis revealed a strong dependence on substrate charge (Figs. 1C and S2). Negatively charged variants (e.g., GFP^−30^–ssrA) were degraded substantially faster than positively charged substrates (e.g., GFP^+15^–ssrA and GFP^+36^–ssrA) (Figs. 1C and S2). To rule out the possibility that substrate charge primarily affected substrate recognition or productive unfolding, we performed Michaelis–Menten analysis over a range of substrate concentrations (Figs. 1D and 1E). *K*^*M*^ values were relatively similar among variants, whereas *V*_*max*_ progressively decreased with increasing positive surface charge. For the most highly charged substrates (GFP^+15^–ssrA and GFP^+36^–ssrA), kinetic parameters could not be reliably determined because degradation was too slow. To determine whether the poor degradation of positively charged substrates resulted from defective activation of the ClpX motor, we measured ATP hydrolysis by ClpXP in the presence of GFP^−7^–ssrA and GFP^+15^–ssrA (Fig. 1F). Both substrates stimulated ATPase activity above the basal level observed for ClpXP alone, although GFP^−7^–ssrA showed a smaller increase than GFP^+15^–ssrA. Thus, the positively charged substrate remains capable of activating the motor despite its inefficient degradation. To exclude the possibility that differences in degradation rates arose from altered protein stability, we measured circular dichroism spectra of each GFP variant from 20– 90°C (Fig. S3). All substrates exhibited comparable secondary structure and thermal stability, indicating that the observed differences in degradation were not attributable to differences in protein stability (Fig. S3), consistent with previous studies showing that these GFP variants possess similar thermal stability^21^.

Next, we asked whether altering the ionic environment would influence degradation kinetics in a manner consistent with electrostatic contributions to substrate processing. Substitution of KCl with NaCl altered both *V*_*max*_ (~2-fold) and *K*_*M*_ (~3-fold) for GFP^−7^–ssrA (Figs. 2A and 2B). Although K^+^ and Na^+^ differ in hydration and ion-pairing properties in addition to electrostatic screening, these results indicate that substrate engagement and unfolding are sensitive to the ionic environment. The observed effects on *K*_*M*_ likely reflect altered interactions between the negatively charged ssrA degron and the positively charged RKH loops involved in the initial substrate recognition^17^, whereas the accompanying changes in *V*_*max*_ suggest that electrostatic interactions also contribute to productive unfolding and/or translocation^17^.

**Figure 2.**
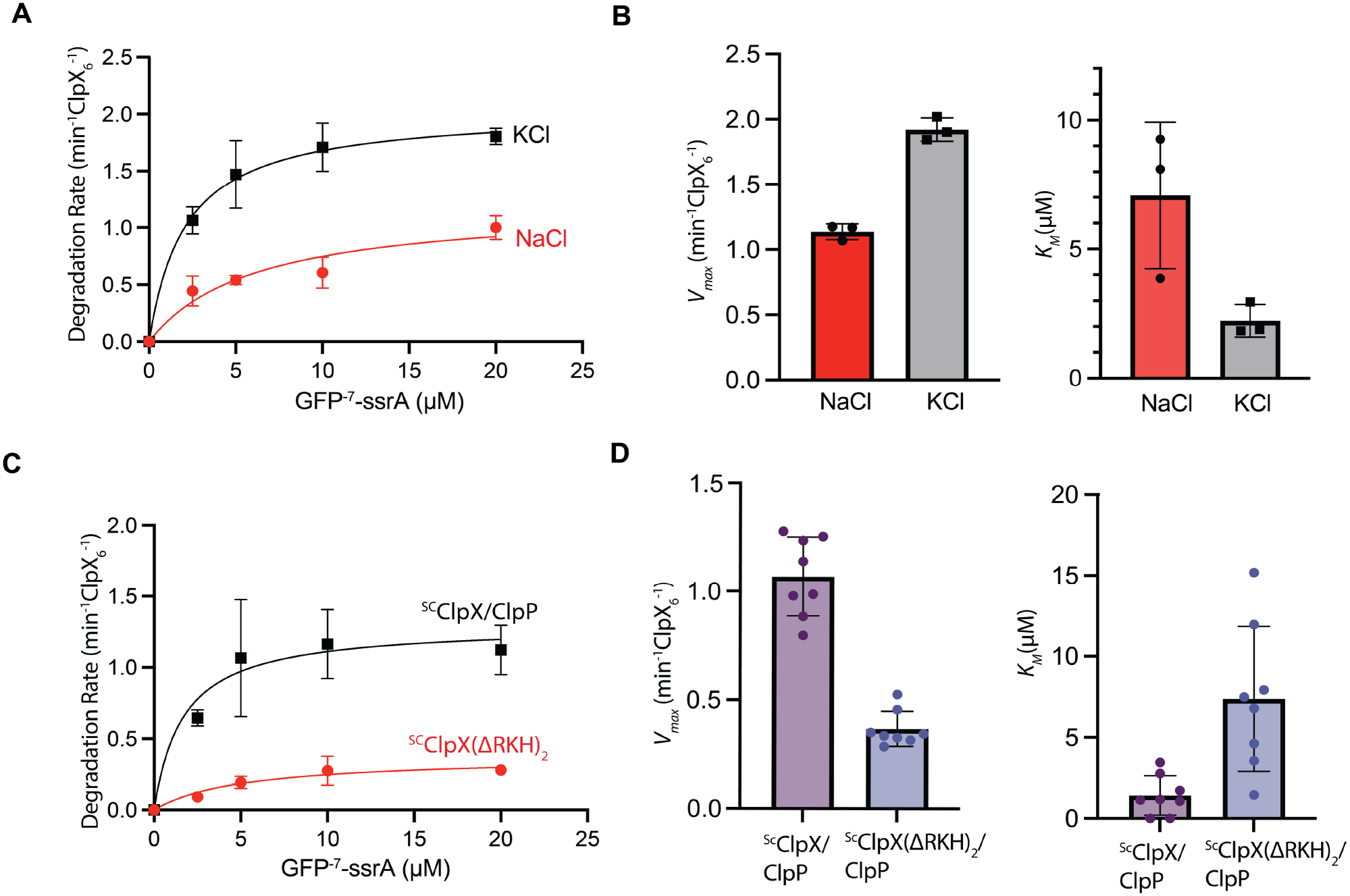
Electrostatic interactions and RKH loops contribute to ClpXP-mediated unfolding. (**A**) The effect of ionic condition (KCl vs. NaCl) on GFP^−7^–ssrA degradation kinetics. (**B**) Comparison of kinetic parameters derived from the fits in panel A. Error bars represent SD from three independent experiments. (**C**) Michaelis–Menten analysis comparing degradation by single-chain ClpX^ΔN^(^sc^ClpX^ΔN^) and a variant lacking two RKH loops in ClpX[^sc^ClpX^ΔN(ΔRKH)2^] with GFP^− 7^–ssrA. Removal of two RKH loops substantially impaired degradation kinetics. Curves represent Michaelis–Menten fits. (**D**) Comparison of kinetic parameters derived from the fits in panel C. Loss of two RKH loops reduced *V*_*max*_ and increased *K*_*M*_. Data points represent independent experiments; bars indicate mean ± SD (n = 7–8 independent experiments).

To further examine the contribution of electrostatic interactions, we varied the KCl concentration while monitoring degradation of negatively and positively charged GFP substrates under conditions of substrate excess (10 µM substrate, 0.5 µM ClpX_6_) (Fig. S4A,B). Increasing ionic strength progressively enhanced degradation of GFP^+15^–ssrA as the KCl concentration was increased from 0 to 400 mM, whereas degradation of GFP^−7^–ssrA decreased primarily at KCl concentrations between 200 and 800 mM. These opposite responses are consistent with electrostatic screening of favorable and unfavorable substrate-ClpX interactions.

#### RKH Loops Directly Enhance the Rate-Limiting Unfolding Step

Complete deletion or alanine substitution of the RKH loops abolishes recognition of ssrA-tagged substrates, preventing separation of their roles in degron binding and mechanical unfolding^19, 22^. To isolate the contribution of the RKH loops to productive substrate denaturation, we employed a previously described single-chain ClpX trimer construct^23^ lacking the N domain [^sc^ClpX^ΔN^], in which deletion of one RKH loop (220-238) per trimer generates a functional hexamer lacking two of the six RKH loops [^sc^ClpX^ΔN(ΔRKH)2^]. Degradation assays using GFP^−7^-ssrA revealed that removal of only two RKH loops impaired substrate recognition, as expected, but also reduced *V*_*max*_ (~3 fold) (Figs. 2C and 2D). Human mitochondrial ClpX contains an RKL loop in place of the bacterial RKH sequence and does not recognize ssrA-tagged substrates^24^. To test whether more conservative changes in loop chemistry alter unfolding, we replaced two bacterial RKH loops with the human mitochondrial RKL sequence [^sc^ClpX^ΔN(ΔRKL)2^]. This substitution increased *K*_*M*_ but did not substantially reduce *V*_*max*_, indicating that the RKL loop largely preserves unfolding capacity while altering degron recognition specificity (Figs. S4C-D).

### Cryo-EM Structures of the ClpXP•GFP^15+^– ssrA Complex Reveal Electrostatically Frustrated Engaged Intermediates

To determine how interactions between the RKH loops and the substrate affect unfolding efficiency, we determined cryo-EM structures of ^sc^ClpX^ΔN^/ClpP complexes engaged with GFP^+15^–ssrA (Figs. S5-S9, and Table S1). Because this positively charged substrate is degraded slowly, we reasoned that substrate-engaged intermediates, in which the folded GFP domain remains positioned at the entrance to the axial channel prior to unfolding, would be sufficiently long-lived to permit structural characterization of the substrate–ClpX interface. Similar substrate-engaged states have previously been observed only under stalled unfolding conditions using methotrexate-bound DHFR–ssrA substrates, in which ligand stabilization renders the substrate highly resistant to unfolding (Fig. S1)^20^. Processing of the ClpXP•GFP^+15^–ssrA dataset yielded three major classes: two substrate-engaged complexes (Figs. S5 and S6) and one class lacking resolved substrate density (Fig. S7). Intermediate I and Intermediate II were reconstructed to overall resolutions of 3.3 and 4.5 Å, respectively, and both contained well-resolved ClpX and ClpP densities together with lower-resolution density corresponding to folded GFP. Nevertheless, density corresponding to an internal GFP α-helix remained visible, enabling reliable docking of an AlphaFold3^25^ model of supercharged GFP and assignment of substrate orientation relative to the ClpX axial channel, particularly after map sharpening with DeepEMhancer^26^. The dominant GFP^+15^–ssrA Intermediate I complex revealed a substrate-engaged conformation in which the folded GFP domain is tilted relative to the ClpX substrate-entry channel, unlike the upright DHFR–ssrA conformation that maximizes contacts with the surrounding RKH loops (Fig. 3A, Fig. S1, and Fig. S9)^20^. Consequently, the tilted GFP orientation positions much of the folded domain away from the RKH ring, restricting most loop–substrate interactions to the narrow entrance of the axial channel (Fig. 3C).

**Figure 3.**
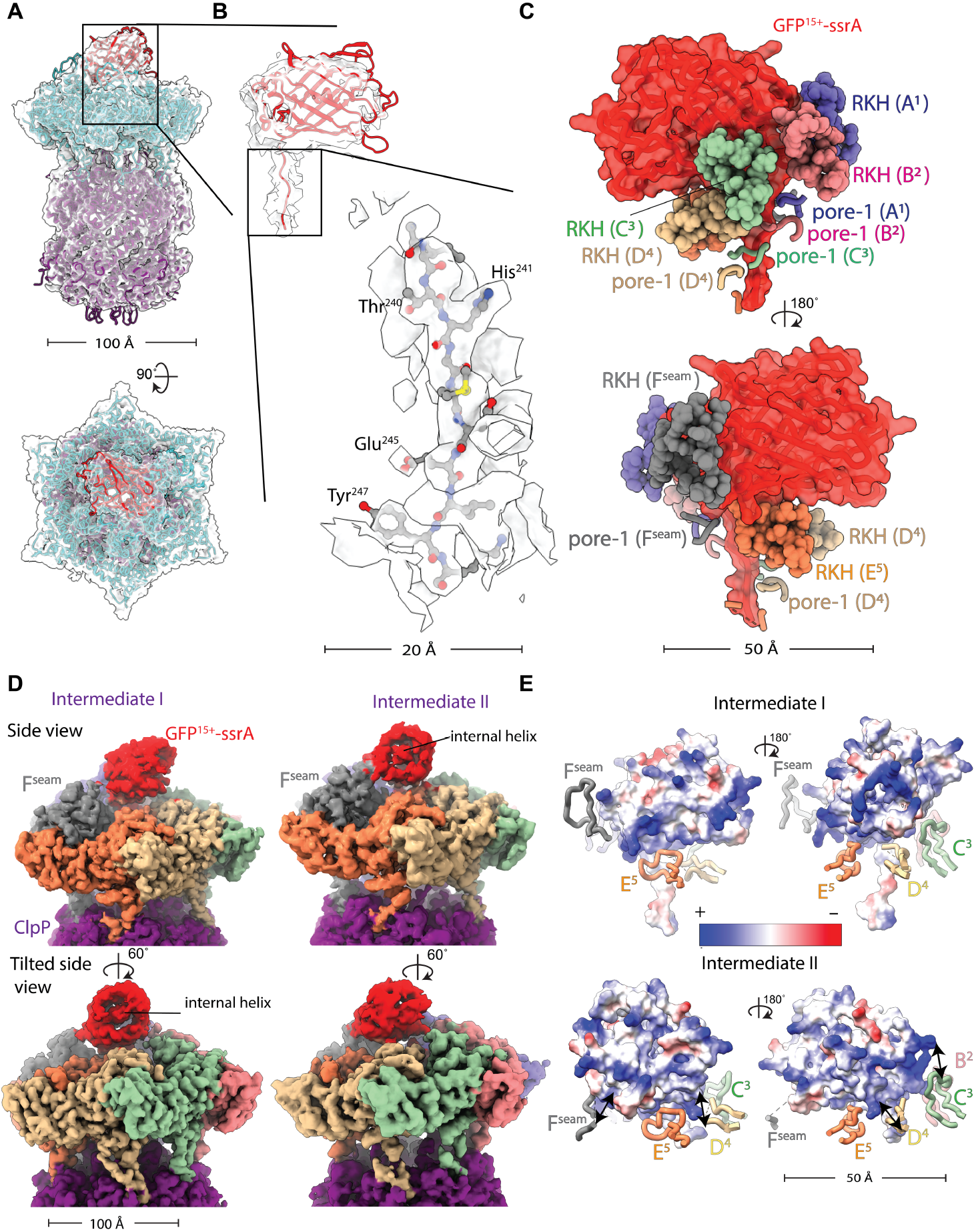
Cryo-EM structures of ^Sc^ClpX^ΔΝ^•ClpP•GFP^+15^–ssrA unfolding intermediates. (**A**) Overall cryo-EM map and atomic model of ClpXP engaged with GFP^+15^–ssrA. ClpX and ClpP is shown as cyan and purple surface representations. The folded GFP^+15^–ssrA domain (red) is engaged at the entrance of the ClpX axial channel. (**B**) Density corresponding to the folded GFP domain and the partially engaged degron is visible within the ClpX axial channel. The inset highlights representative side-chain density for substrate residues within the ClpX axial channel. (**C**) The folded portion of GFP^+15^–ssrA interacts with RKH loops (space filling representation) and pore-1 loops (cartoon representation) from multiple ClpX subunits. Notably, the pore-1 loops adopt a spiral arrangement around the threaded region of the substrate, consistent with a translocation-competent conformation. (**D**) Two distinct GFP^+15^–ssrA substrate-engaged intermediates identified by cryo-EM 3D classification. Relative to Intermediate I (panels A–C), Intermediate II shows a rotated orientation of the folded GFP domain toward the ClpX seam subunit, as indicated by the position of a resolved internal GFP α-helix. Cryo-EM maps are colored according to the corresponding atomic models. Side and tilted views reveal distinct orientations of the folded substrate domain relative to the ClpX axial channel. The maps were sharpened using DeepEMhancer^26^. (**E**) Electrostatic surface representations of the two GFP^+15^–ssrA intermediates. In both conformations, positively charged regions of the substrate are positioned adjacent to the positively charged RKH loops at the channel entrance. Relative to Intermediate I, Intermediate II presents a larger positively charged surface of GFP^+15^ toward the RKH ring, as indicated by the double arrows, consistent with a more electrostatically unfavorable interface.

The density corresponding to the threaded substrate within the ClpX axial channel was resolved at higher resolution, facilitating substrate modeling and register assignment (Figs. 3A and 3B). Consistent with previously reported substrate-engaged ClpX structures, the pore-1 loops adopted the canonical spiral arrangement around the threaded substrate that is characteristic of ATP-driven substrate translocation ^12^ (Fig. 3C). Despite adopting this apparently productive substrate-engaged configuration, GFP^+15^–ssrA was degraded inefficiently (Figs. 1 and S2), indicating that substrate engagement alone is insufficient to account for fruitful unfolding. Electrostatic surface analysis further revealed that the region of GFP^+15^ positioned adjacent to the substrate-entry channel is enriched in positive charge, consistent with an electrostatically unfavorable interface with the positively charged RKH loops (Fig. 3E). Consistent with this interpretation, several RKH loops exhibited weaker density, suggesting heterogeneous loop– substrate interactions that may alleviate these unfavorable electrostatic contacts. Structural overlay of the GFP^+15^–ssrA and DHFR-engaged complexes revealed that the principal conformational differences were localized to the RKH loops (Fig. S10), consistent with the idea that electrostatic incompatibility at the substrate-entry channel perturbs productive loop–substrate interactions during unfolding.

A second, less populated GFP^+15^–ssrA Intermediate II class revealed an additional rotation of the folded GFP domain relative to Intermediate I (Fig. 3D). Although the overall resolution of Intermediate II (Fig. S7) was lower than that of Intermediate I (Fig. S6), an internal GFP α-helix remained sufficiently well resolved to support assignment of a distinct substrate-engagement orientation. Notably, this orientation positions a larger positively charged surface of GFP adjacent to the RKH ring, consistent with a more electrostatically unfavorable interface (Fig. 3E, bottom panel).

Inspired by these substrate-engaged intermediates, we next asked whether GFP^+15^–ssrA remains persistently bound to ClpX, thereby blocking access of competing substrates and inhibiting their degradation. To address this question, we preincubated GFP^+15^–ssrA with ClpXP before adding Arc–ssrA as a competing ssrA-tagged substrate. Arc–ssrA was degraded rapidly in the absence of GFP^+15^–ssrA preincubation, whereas prior incubation with GFP^+15^–ssrA delayed, but did not prevent, Arc–ssrA degradation (Fig. S11). These results indicate that GFP^+15^–ssrA does not remain persistently trapped within the ClpX substrate-entry channel and is eventually released. Consistent with this interpretation, focused 3D classification of the GFP^+15^–ssrA dataset identified a third class (intermediate III) lacking resolved substrate density (Fig. S8 and Table S1), likely representing complexes following substrate disengagement from the axial channel.

### RKH-Loop Deletion Accumulates Substrate-Engaged Unfolding Intermediates

To further examine how the RKH loops contribute to productive unfolding, we analyzed cryo-EM structures of the negatively charged substrate GFP^−7^–ssrA complexed by the partially loop-deficient enzyme [^sc^ClpX^ΔN(ΔRKH)2^/ClpP] (Figs. S12-15, Fig. 4, and Table S1). Because GFP^−7^–ssrA is efficiently degraded by ClpXP, we reasoned that combining this substrate with the slowly degrading ΔRKH ClpXP enzyme would prolong transient substrate-engaged unfolding intermediates that are normally too short-lived to visualize. Focused 3D classification of the GFP^−7^–ssrA dataset revealed multiple substrate-engaged conformations containing distinct substrate densities positioned above the ClpX axial channel (Fig. 4A and Figs. S12–S15). Several classes refined to global resolutions of ~2.9–3.6 Å, yielding well-resolved density for the ClpX and ClpP assemblies (Fig. 4). Despite substantial conformational heterogeneity, density corresponding to the folded GFP domain was resolved in several maps representing ~50% of particles (Fig. 4A). However, only one class (Map a; ~19.5% of particles) contained sufficiently resolved density for an internal GFP α-helix and the threaded degron to enable confident assignment of substrate orientation and register within the axial channel (Fig. S13).

**Figure 4.**
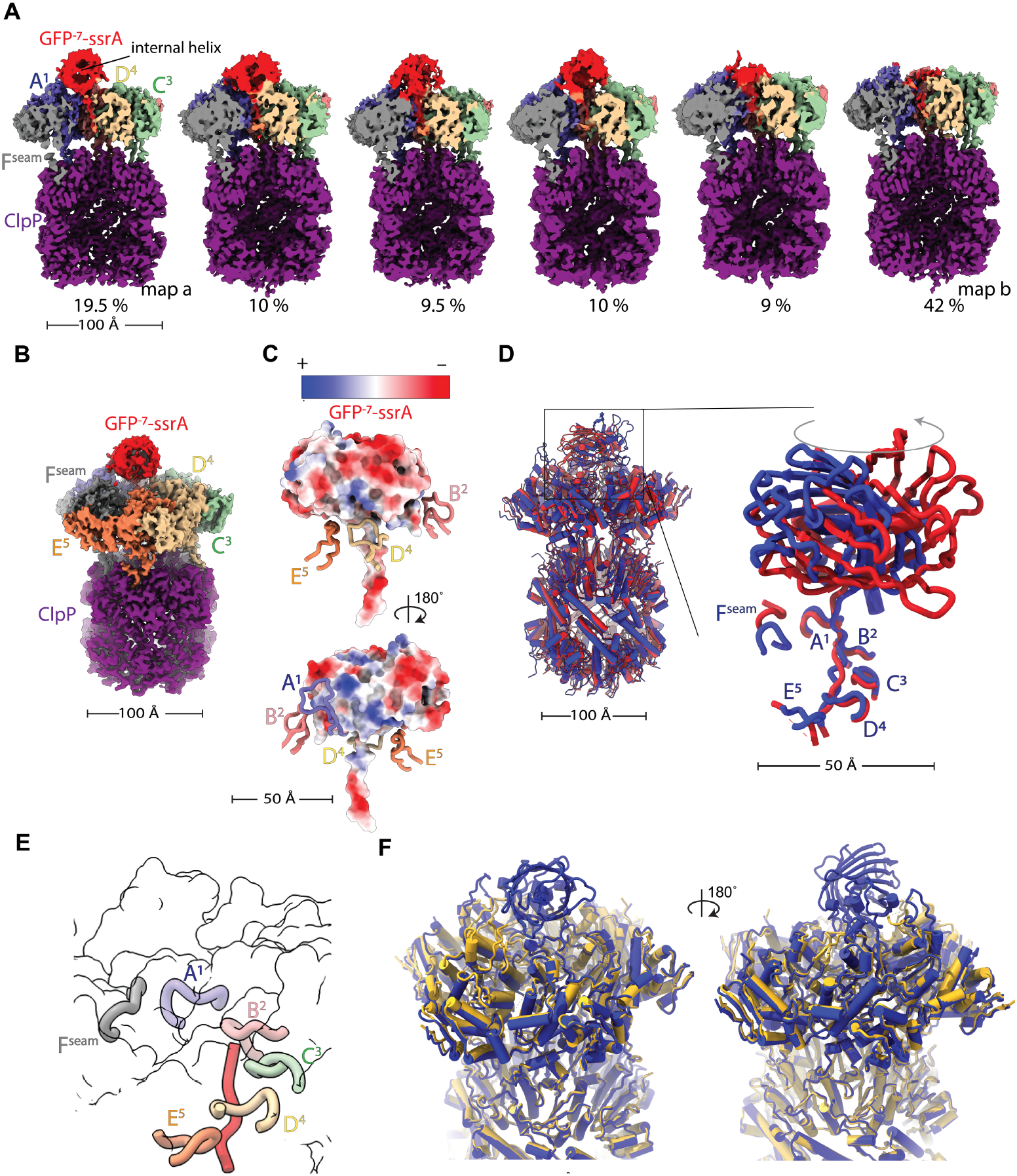
Cryo-EM structures of ^sc^ClpX^ΔN(ΔRKH)2^•ClpP•GFP^−7^–ssrA engaged by an RKH-loop deletion mutant of ClpX. (**A**) Cryo-EM maps corresponding to multiple substrate-engaged conformations identified by focused 3D classification of the GFP^−7^–ssrA•[^sc^ClpX^ΔN(ΔRKH)2^]•ClpP complex. Maps are colored according to the fitted atomic models. Individual ClpX subunits are colored separately, whereas ClpP is shown in purple. Percentages indicate the fraction of particles corresponding to each reconstructed class. (**B**) Cryo-EM map of the GFP^−7^–ssrA-engaged state (19.5% particle class, Map a) bound to the RKH-loop deletion mutant [^sc^ClpX^ΔN(ΔRKH)2^]**•**ClpP. Cryo-EM density is colored according to the corresponding atomic model. ClpP is shown in purple, ClpX subunits are colored individually, and GFP^−7^–ssrA is shown in red. (**C**) Electrostatic surface representation of the substrate-engaged complex shown in panel B. Color-coded RKH loops from individual ClpX subunits surround the folded GFP^−7^–ssrA substrate at the entrance of the axial channel, positioning positively charged RKH loop residues adjacent to negatively charged regions of the substrate. (**D**) Structural comparison of GFP^−7^–ssrA engaged by [^sc^ClpX^ΔN(ΔRKH)2^] **•**ClpP (blue) and GFP^+15^–ssrA Intermediate I (red). Relative to GFP^+15^–ssrA Intermediate I, the negatively charged GFP^−7^ substrate adopts a rotated orientation that positions a more negatively charged surface toward the substrate-entry channel. (**E**) Atomic model corresponding to the 42% particle class (Map b), showing an unidentified segment of substrate threaded through the ClpX axial channel, consistent with previously characterized translocation complexes^12^. The substrate density was modeled as polyalanine. Pore-1 loops are shown in cartoon representation and colored according to ClpX subunit. (**F**) Structural comparison of the fully engaged GFP^−7^–ssrA intermediate (blue; Map a) and the translocation complex (yellow; Map b), corresponding to the 19.5% and 42% particle populations shown in panel A, respectively. The overall ClpX conformations are highly similar, despite substantial remodeling of the substrate.

In contrast to the dominant GFP^+15^–ssrA Intermediate I state, the folded GFP^−7^ domain adopted a rotated orientation that more closely resembled the GFP^+15^–ssrA Intermediate II conformation (Fig. 4D). Electrostatic surface analysis indicated that this orientation positions negatively charged regions of the substrate adjacent to the positively charged RKH loops (Fig. 4C), in contrast to the electrostatically unfavorable interfaces observed in the GFP^+15^–ssrA structures (Fig. 3).The most abundant class (~42% of particles) displayed a conformation resembling previously described ClpXP translocation complexes^12, 20^, with an unidentified substrate density extending through the axial channel and modeled as poly-alanine because the density was insufficient to resolve side-chain features (Fig. 4E). The pore-1 loops retained the characteristic helical arrangement around the translocating substrate (Fig. 4E), consistent with numerous previously reported AAA protease translocation intermediates^17, 27^.

Although we refrain from making definitive structural assignments for the additional classes identified by focused classification (Fig. 4A), the presence of one well-resolved folded GFP-engaged state (Fig. 4B), together with multiple maps exhibiting varying extents of GFP density above the ClpX channel, suggests that these structures may represent partially unfolded substrate intermediates captured during denaturation attempts. To further evaluate the structural heterogeneity of our complex, we performed cryoDRGN analysis^28–29^, which revealed variability in substrate position and density relative to the ClpX axial channel that was consistent with the focused 3D classification results, further supporting the existence of multiple substrate conformational states (Movie 1).

Structural overlay of the translocation-like complex (Map b) with the folded substrate-engaged state (Map a) revealed that substantial substrate remodeling can occur without major rearrangement of the overall ClpX spiral conformation (Fig. 4F). Similar observations were previously reported for comparisons between the methotrexate-bound DHFR–ssrA substrate-engaged complex (PDB ID: 8V9R) and translocation complexes containing unfolded DHFR– ssrA (PDB ID: 9C88)^20^, suggesting that ClpXP-mediated unfolding may generally proceed through substrate remodeling within a relatively conserved motor conformation.

## DISCUSSION

AAA+ proteases must mechanically unwind the stable native structures of proteins before degradation, making unfolding an essential step in proteolysis^3, 20, 30^. Failure of this process is particularly detrimental because substrates can become partially processed, lose their recognition degrons, and escape further degradation while accumulating in the cell. Efficient unfolding of the diverse substrates generated during translation and cellular stress^11^ therefore represents a major challenge for AAA+ proteases.

Here, using biochemical assays with a panel of supercharged GFP variants bearing the same degron sequence and exhibiting comparable thermal stability, together with cryo-EM structures, we identify electrostatic complementarity between the positively charged entrance of the ClpX substrate-entry channel and folded substrate domains as a key determinant of productive ATP-driven substrate unfolding. Our cryoEM structures show that the positively charged GFP^+15^–ssrA populates multiple substrate-engaged orientations (Figs. 3D and 3E), likely reflecting repeated attempts to optimize interactions with the positively charged RKH loops. We propose that these complexes represent electrostatically frustrated states in which the enzyme samples alternative substrate orientations to alleviate unfavorable surface-charge interactions (Fig. 5, top panel). Notably, these states retain proper pore-loop engagement with the threaded region of the substrate, and GFP^+15^–ssrA stimulates ATP hydrolysis above basal ClpXP levels (Fig. 1F), indicating that degron engagement, mechanical grip, and motor activation alone are insufficient to drive efficient substrate unfolding. Together, these observations and the opposite effects of increasing ionic strength on GFP^−7^–ssrA and GFP^+15^–ssrA degradation rates (Fig. S4A,B) support the conclusion that the observed effects arise from electrostatic interactions.

**Figure 5.**
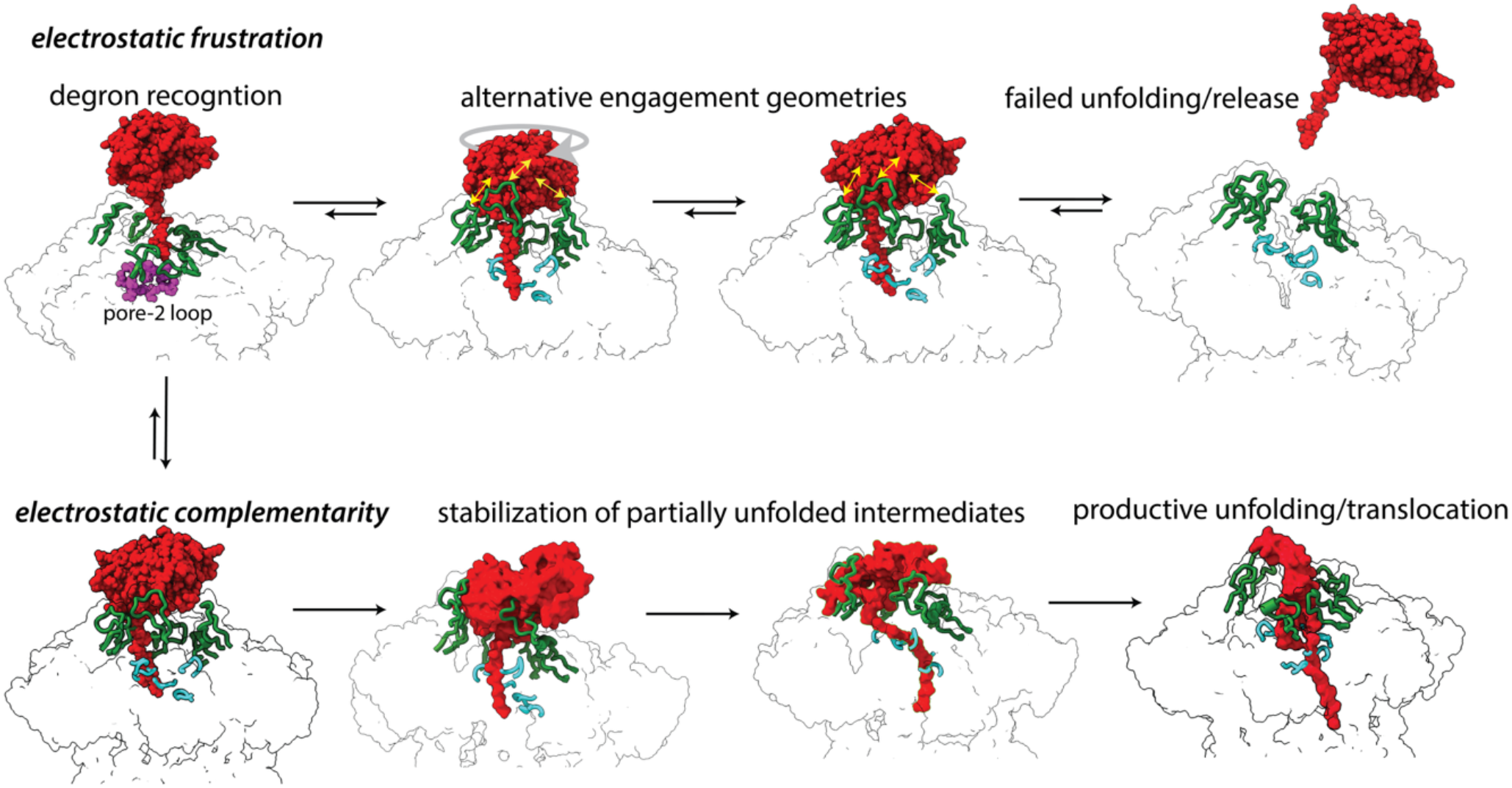
Proposed model for electrostatic regulation of substrate unfolding by ClpXP. **Upper pathway**: Following substrate recognition, translocation and engagement, positively charged substrate surfaces generate electrostatic frustration at the ClpX channel entrance through unfavorable interactions with the positively charged RKH loops (green). These interactions promote repeated sampling of alternative substrate-engagement geometries, which may ultimately result in substrate release. **Lower pathway:** Favorable electrostatic complementarity between the substrate surface and the RKH loops stabilizes transient, partially unfolded intermediates generated during successive ATP-driven unfolding attempts, thereby reducing substrate backsliding and promoting productive unfolding and translocation. Partially unfolded substrate conformations are shown schematically and do not represent independently refined atomic models. The substrate is shown in red, the RKH loops in green, the pore-1 loops in cyan, and the pore-2 loop of chain A in pink (sphere representation). ClpP is omitted for clarity. Yellow arrows indicate electrostatic repulsion between positively charged substrate surfaces and the positively charged RKH loops.

Biochemical experiments using single-chain ClpX lacking two RKH loops (Figs. 2C and 2D), together with the corresponding cryo-EM structures (Fig. 4), suggest that the RKH loops contribute not only to the early steps of degron recognition, as previously shown^17^, but also to the stabilization of unfolding intermediates during repeated ATP-driven unfolding attempts (Fig. 5). The presence of multiple substrate-engagement states in GFP^−7^–ssrA complexes assembled with RKH-deficient ClpX (Fig. 4A, Movie1) further suggests that ClpXP-mediated unfolding proceeds through progressive, step-wise destabilization of substrate structure rather than cooperative global denaturation analogous to chemical unfolding^31^. This interpretation is consistent with previous single-molecule spectroscopy and optical trapping studies indicating that the ClpXP AAA+ protease unfolds substrates through multiple successive pulling and partial-unfolding events rather than a single denaturation step^32^. In this model, the RKH loops may stabilize transiently unfolded conformations generated during repeated ATP-driven unfolding attempts. Alternatively, by forming contacts with partially unfolded substrate regions, the RKH loops improve mechanical force transmission and limit substrate backsliding, thereby promoting productive protein unfolding (Fig. 5, bottom panel).

However, if positively charged RKH loops can impair unfolding of positively charged substrates, why has ClpX evolved a positively charged substrate-entry channel? One possibility is that the RKH loops represent an adaptation for processing the predominantly negatively charged bacterial cytosolic proteome, whereas highly positively charged proteins may inherently be poor ClpXP substrates and therefore constitute only a minor fraction of physiological targets. Much of the *E. coli* cytosolic proteome carries a net negative charge, a property thought to promote efficient diffusion by minimizing interactions with negatively charged ribosomes^33^. Notably, the RKH motif is conserved specifically within the ClpX family of AAA+ unfoldases^22^, whereas other AAA+ proteases contain distinct substrate-entry elements with different properties. Human mitochondrial ClpX, for example, contains an RKL loop in place of the bacterial RKH motif and does not efficiently recognize ssrA-tagged substrates^27^. Recent studies have proposed that human ClpXP recognizes phosphorylated substrates in which phosphorylation functions as a degron^24^. Because phosphorylation introduces negative charge, our findings raise the possibility that favorable electrostatic interactions with phosphorylated substrates may also promote productive substrate engagement and degradation by hClpXP. More broadly, differences in substrate-entry channel chemistry may help explain why distinct AAA+ proteases–including ClpXP, ClpAP, HslUV, and FtsH–can recognize similar ssrA degron sequences yet display different substrate preferences and unfolding efficiencies^34–35^. For example, the more hydrophobic environment of the FtsH substrate-entry channel may be better suited for membrane-protein quality control and degradation of hydrophobic membrane-bound ssrA-tagged substrates^36–38^.

Modulation of ClpXP activity has recently emerged as a promising strategy in both human disease and bacterial infection, including inhibition of mitochondrial ClpXP in certain cancers^8, 39^ and targeting of ClpXP-dependent proteolysis in bacterial pathogens such as *Mycobacterium tuberculosis*^6, 40^. Our results suggest that perturbation of substrate-entry electrostatics could represent a previously unrecognized mechanism for modulating ClpXP activity, either by destabilizing productive unfolding intermediates or by trapping nonproductive substrate-engaged states. Together, our results demonstrate that productive unfolding by AAA+ proteases depends not only on degron recognition and ATP-driven force generation, but also on electrostatic complementarity at the substrate-entry channel.

## Materials and Methods

### Plasmid Construction and Protein Expression

GFP charge variants GFP^−7^ and GFP^+7^ were generously provided by the Poolman laboratory (Schavemaker *et al*., eLife 2017), and GFP^−30^ was obtained from Addgene (Addgene #199167). All GFP variants were cloned into a pETDuet expression vector under a T7 promoter and contain an N-terminal His_6_-tag (AmpR). The C-terminal ssrA sequence (AANDENYALAA) was appended to each construct by PCR and Gibson Assembly. All ClpX constructs used in this study are N-terminal domain–deleted ClpX (ClpX^ΔN^). The ΔRKH variant was generated by deleting residues 220–238 in the monomeric ClpX sequence using Q5 Site-Directed Mutagenesis (NEB) and KLD recircularization. Likewise, the RKL variant was generated by introducing the H230L mutation in monomeric ClpX using the same mutagenesis workflow. Each verified monomeric mutant was then substituted into subunit A of the single-chain ClpX using Gibson Assembly, leaving the remaining subunits unchanged.

### Protein expression and purification

Proteins were expressed in *E. coli* T7 *ΔclpX/ΔclpP/ΔclpA* cells transformed with plasmids encoding His_6_-tagged GFP variants or ClpX/ClpP. For ClpX and GFP variants, cells were grown in LB medium containing the appropriate antibiotic at 37°C until reaching an OD_600_ of 0.6–0.8, induced with 0.5 mM IPTG, and incubated at 16°C overnight. Cells were harvested by centrifugation at 4000 rpm for 25 min at 4°C and resuspended in 10 mL lysis buffer [20 mM HEPES pH 7.5, 400 mM NaCl, 100 mM KCl, 10% glycerol, 1 mM DTT, 20 mM imidazole, and protease inhibitor tablet] per liter of culture. Cells were lysed by sonication on ice, and the lysate was clarified by centrifugation at 14,000 rpm for 30 min at 4°C. The supernatant was loaded onto a Ni^2+^-NTA column pre-equilibrated with the same buffer, and eluted with buffer containing 250 mM imidazole. Eluted fractions containing the target protein were pooled, concentrated, and further purified by size-exclusion chromatography using a Superdex 200 column equilibrated with 20 mM HEPES pH 7.5, 300 mM KCl, 10% glycerol, and 1 mM DTT. ClpP was expressed in LB medium supplemented with the appropriate antibiotic. Cells were grown at 37°C to an OD_600_ of 0.6–0.8, induced with 0.5 mM IPTG, and incubated with shaking at 30°C for 3 h. Cells were harvested by centrifugation at 4,000 rpm for 25 min at 4°C and resuspended in 10 mL of ClpP lysis buffer per liter of culture (50 mM sodium phosphate, pH 8.0, 1 M NaCl, 10% glycerol, 5 mM imidazole, and 1 mM DTT). Cells were lysed by sonication on ice, and the lysate was clarified by centrifugation at 14,000 rpm for 30 min at 4°C. The supernatant was loaded onto a Ni^2+^-NTA column pre-equilibrated with the same buffer, washed with buffer containing 40mM imidazole, and eluted with buffer containing 500 mM imidazole. The protein was further purified by size-exclusion chromatography using a Superdex 200 column equilibrated with 50 mM Tris–HCl pH 8.0, 150 mM KCl, 10% glycerol, 0.5 mM EDTA, and 1 mM DTT. Purified proteins were concentrated, flash-frozen in liquid nitrogen, and stored at −80 °C.

### GFP degradation assay

GFP–ssrA degradation was monitored by measuring the decrease in fluorescence (excitation 467 nm; emission 511 nm) using a SpectraMax M2 plate reader. Concentrations of ClpX variants, ClpP and substrates were measured using a NanoDrop spectrometer (ThermoFischer Scientific) at an absorbance of 280 nm, and concentrations were calculated using extinction coefficients obtained from https://web.expasy.org/protparam. Reactions were performed at 30°C in buffer containing 25 mM HEPES (pH 7.5), 5 mM MgCl_2_, 200 mM KCl, and 10% glycerol, supplemented with 5 mM ATP, 32 mM creatine phosphate, and 0.08 mg/mL creatine kinase. Each reaction contained 0.5 μM ClpX_6_ (or variants) and 1.5 μM ClpP_14_. GFP–ssrA substrate of various concentrations to determine kinetic parameters. All components except substrate were preincubated in 384-well assay plate (Corning, 3575) at 30 °C for 5 min, and reactions were initiated by adding pre-equilibrated substrate. Fluorescence intensity was recorded by SpectraMax M2 plate reader, and initial degradation rates were determined from the linear portion of the fluorescence decay curve. Michaelis–Menten parameters (*K*_*M*_ and *V*_*max*_) were obtained by fitting the initial rates as a function of substrate concentration using GraphPad Prism. NaCl dependence was tested by performing the degradation assay in identical buffer conditions but substituting 200 mM KCl with 200 mM NaCl, while keeping all other components constant. Gel-based degradation assays were performed under the same reaction conditions as the fluorescence assay, except that reactions were quenched at the indicated time points by the addition of Coomassie Blue SDS loading buffer and heated at 95 °C for 5 min. Samples were resolved on 4–20% Mini-PROTEAN TGX gels (Bio-Rad) and imaged using a Bio-Rad imaging system.

### ATPase activity assay

ATP hydrolysis was monitored using a coupled enzymatic assay as described by Norby (1988), in which oxidation of NADH to NAD^+^ was measured as a decrease in absorbance at 340 nm (Δε = 6.22 mM^−1^ cm^−1^). Absorbance changes were recorded using a Synergy HTX multimode plate reader in 384-well assay plates (Corning 3575).

A 20x coupling reagent was prepared by combining pyruvate kinase/lactate dehydrogenase from rabbit muscle (Sigma-Aldrich, P0294), NADH, and phosphoenolpyruvate in 25 mM HEPES-KOH (pH 7.6). ATPase reactions were assembled by mixing equal volumes of protein solution and a 2X ATPase master mix. Final reactions contained 0.5 µM ClpX_6_, 1.5 µM ClpP_14_, 5 µM GFP^+15^–ssrA or GFP^−7^–ssrA, and 5 mM ATP. Control reactions contained ClpXP in the absence of substrate. Reactions were initiated by addition of ATP, and ATP hydrolysis rates were determined from the linear decrease in absorbance at 340 nm using the extinction coefficient of NADH.

### Circular Dichroism Spectroscopy

Circular dichroism (CD) spectra were obtained by a Chirascan spectropolarimeter (Applied Photophysics). Proteins were diluted to a final concentration of 5 µM in phosphate-buffered saline (PBS) for analysis. Measurements were performed in a 10 mm pathlength quartz cuvette. Thermal denaturation was monitored by collecting full far-UV CD spectra from 200 to 280 nm as the temperature was ramped from 20 to 90°C at a rate of 1°C min^−1^. Spectra were acquired with a bandwidth of 1 nm, a step size of 1 nm, and a time per point of 0.5 s; each reported spectrum represents an average of 3 accumulations.

### Cryo-EM sample preparation

For GFP^+15^–ssrA/ClpXP structure, purified ^SC^ClpX^ΔN^ and ClpP complexes were mixed to final concentrations of 5.7 µM ^SC^ClpX_6_^ΔN^ and 1.5 µM ClpP_14_ in 20 mM HEPES (pH 7.5), 100 mM KCl, and 25 mM MgCl_2_, and incubated with 5 mM ATP for 5 min at room temperature. GFP^15+^–ssrA substrate (20 µM) was then added to initiate the reaction. A 2.5 µL aliquot of the reaction sample was applied to Quantifoil R2/1 Cu 200 mesh grids that had been glow-discharged for 60 s at 25 mA using a GloQube Plus (MiTeGen) 3 minutes after the initiation of reaction. Grids were blotted for 4 s with a blot force of +4 at 6°C and 100% relative humidity in a FEI Vitrobot Mark IV instrument (Thermo Fisher Scientific).

For ^sc^ClpX^ΔN(ΔRKH)2^ ClpXP structure bound with GFP^−7^–ssrA, 5.7 µM ^SC^ClpX_6_^ΔN(ΔRKH)2^ and 1.5 µM ClpP_14_ were mixed in 20 mM HEPES (pH 7.5), 100 mM KCl, and 25 mM MgCl_2_, and incubated with 5 mM ATP for 5 min at room temperature. GFP^−7^-ssrA substrate (20 µM) was then added to initiate the reaction. A 2.5 µL aliquot of the reaction sample was applied to Quantifoil R2/1 Cu 200 mesh grids that had been glow-discharged for 60 s at 25 mA using a GloQube Plus (MiTeGen) at 5 minutes after the initiation of reaction. Grids were blotted for 4 s with a blot force of +4 at 6°C and 100% relative humidity in a FEI Vitrobot Mark IV instrument (Thermo Fisher Scientific).

### Cryo-EM data collection

For the GFP^+15^–ssrA structure, two independent datasets were collected using EPU on a Titan Krios G3 operated at an acceleration voltage of 300 kV and equipped with a Falcon 4 detector at a nominal magnification of 75,000x, corresponding to a calibrated pixel size of 0.87 Å/pixel. The first dataset consisted of 6,238 movies and the second dataset consisted of 4,634 movies, resulting in a combined total of 10,872 movies. Movies were recorded with a total accumulated electron exposure of 51.8 e^−^/Å^2^ and a defocus range of −0.4 to −2.0 µm. For the GFP^−7^–ssrA dataset, 5,998 movies were collected using EPU on a Titan Krios G3 operated at 300 kV and equipped with a Falcon 4 detector at a nominal magnification of 130,000x, corresponding to a calibrated pixel size of 0.776 Å/pixel. Movies were recorded with a total accumulated electron exposure of 55.48 e^−^/Å^2^ and a defocus range of −0.4 to −2.0 µm.

### Cryo-EM pre-processing and particle picking

For the GFP^+15^ –ssrA structure, the two datasets were initially processed independently in cryoSPARC using default parameters unless otherwise specified.

For the first dataset, raw movies (6,238) were pre-processed using ‘Patch Motion Correction’ followed by ‘Patch CTF Estimation’. Initial particles were identified using Blob Picker with a circular diameter range of 170–250 Å, yielding 467,127 particles. Particles were extracted using a 440-pixel box size Fourier-cropped to 256 pixels and subjected to 2D classification. Selected 2D classes were subsequently used for ‘Template Picker’, yielding 2,945,341 particles that were extracted with a 440-pixel box size Fourier-cropped to 128 pixels. After multiple rounds of 2D classification, 119,254 particles were retained. An initial ‘*ab initio* reconstruction’ using two classes identified one class corresponding to the target complex containing 47,811 particles. These particles were re-extracted with a 416-pixel box size Fourier-cropped to 256 pixels and subjected to a second round of *ab initio* reconstruction followed by homogeneous refinement, yielding 47,743 particles. To further improve particle quality, ‘Topaz training’ and extraction were performed using an expected particle diameter of 250 Å. Newly extracted particles were subjected to additional rounds of 2D classification, resulting in 151,745 selected particles.

For the second dataset of GFP^+15^-ssrA, raw movies (4,634) were pre-processed using ‘Patch Motion Correction’ and ‘Patch CTF Estimation’. Initial particles were identified using ‘Blob Picker’ with a circular diameter range of 170– 250 Å, yielding 617,101 particles. Particles were extracted using a 440-pixel box size Fourier-cropped to 256 pixels and subjected to 2D classification. Selected 2D classes were used for ‘Template Picker’, and extracted particles (440-pixel box size Fourier-cropped to 128 pixels) were subjected to additional rounds of 2D classification, resulting in 89,383 selected particles. To further improve particle picking, ‘Topaz training’ and extraction were performed using an expected particle diameter of 250 Å. Newly extracted particles were subjected to further rounds of 2D classification, producing 112,274 selected particles.

For the GFP^−7^–ssrA dataset, particles were initially identified using ‘Blob Picker’ and extracted with a box size of 440 pixels, Fourier-cropped to 256 pixels. Following 2D classification, two representative classes corresponding to side and tilted-side views were selected as templates for Template Picker (particle diameter = 180 Å). Template-picked particles were extracted with a box size of 440 pixels, Fourier-cropped to 128 pixels, and subjected to two additional rounds of 2D classification, yielding a preliminary particle stack containing 235,351 particles.

### Ab initio reconstruction and global refinement

For the second GFP^+15^–ssrA dataset, particles were subjected to ‘ab initio reconstruction’ into two classes, from which the class corresponding to the ClpXP complex was selected. Particles from this class were re-extracted using a 440-pixel box size Fourier-cropped to 284 pixels and subjected to a second round of ‘ab initio reconstruction’ followed by ‘homogeneous refinement’. Particles from both datasets were subsequently merged for downstream refinement and conformational analysis. The merged particle stack, containing 112,258 particles, was subjected to homogeneous refinement. A soft mask encompassing the substrate was generated in ChimeraX and used for subsequent 3D classification in cryoSPARC, yielding eight classes. The eight classes were subsequently combined using Regroup 3D Classes (3 classes).

One class, containing 76,894 particles, was subjected to ‘homogeneous refinement’ followed by ‘Global CTF Refinement’ and ‘non-uniform refinement’. Further particle alignment using ‘Volume Alignment Tools’ to center on ClpX and substrate and subsequent ‘local refinement’ improved map quality around the substrate. Final reconstructions were generated through iterative rounds of ‘non-uniform refinement’, yielding a map with a resolution of 3.3 Å according to the gold-standard FSC criterion. The final map was further enhanced using ‘DeepEMhancer’^26^. Model building was performed using a combination of ChimeraX-1.3 ^41^, Coot-0.9.4 ^42^, and Phenix-1.21-5190 ^43^. Briefly, the AlphaFold3 model of GFP^+15^– ssrA together with the previously determined ClpXP substrate-engaged structure (PDB ID: 8V9R) were docked into the cryo-EM density using ChimeraX. The resulting model was manually adjusted in Coot and subsequently refined using real-space refinement in Phenix. Multiple rounds of manual rebuilding and automated refinement were performed until convergence of model geometry and map fit.

A second subset containing 29,496 particles was selected for non-uniform refinement, followed by ‘Global CTF Refinement’ and an additional round of ‘non-uniform refinement’. Further particle alignment was performed using ‘Volume Alignment Tools’, followed by ‘non-uniform refinement’, resulting in a map with a resolution of 4.5 Å according to the gold-standard FSC criterion. The final map was further enhanced using DeepEMhancer ^26^.

A third class containing 19,195 particles was subjected to ‘homogeneous refinement’, ‘Global and Local CTF Refinements’, and subsequent rounds of ‘non-uniform refinement’ and ‘local refinement’, yielding Intermediate III at a resolution of 4.26 Å according to the gold-standard FSC criterion.

For the GFP^−7^–ssrA dataset, 235,351 particles were subjected to ‘ab initio reconstruction’ using two classes. One class corresponding to the ClpXP complex, containing 172,926 particles, was selected and re-extracted using a 440-pixel box size Fourier-cropped to 128 pixels, followed by ‘ab initio reconstruction’. Particles were subsequently re-extracted using a 440-pixel box size Fourier-cropped to 320 pixels and subjected to a second round of ‘ab initio reconstruction’.

A soft mask encompassing ClpX and the cis ClpP ring was generated in ChimeraX and used together with ‘Volume Alignment Tools’ to center particles relative to the masked region. The aligned particles were subjected to ‘homogeneous refinement’, and the resulting reconstruction was used for subsequent 3D classification in cryoSPARC, yielding ten classes. The ten classes were subsequently combined using ‘Regroup 3D Classes’ and further classified into six classes.

Two particle subsets containing 33,469 and 71,818 particles were selected for downstream refinement. For ‘Map a’, the 33,469-particle subset was refined using local refinement, yielding a reconstruction with a resolution of 3.5 Å according to the gold-standard FSC criterion. For ‘Map b’, the 71,818-particle subset was subjected to heterogeneous refinement using two classes. A class containing 48,607 particles was selected and further refined by homogeneous refinement, yielding a reconstruction with a global resolution of 2.9 Å according to the gold-standard FSC criterion. Model building was performed using a combination of ChimeraX-1.3 ^41^, Coot-0.9.4 ^42^, and Phenix-1.21-5190 ^43^. For the model building into map a similar protocol was used as described for intermediate I GFP^+15^–ssrA, for map b (PDB ID: 9C88) was used as initial model docked using ChimeraX, with manual model building using coot, and phenix.

### CryoDRGN analysis

To investigate structural heterogeneity, we analyzed all 171,179 particles from the GFP^−7^–ssrA dataset using CryoDRGN v2.3.0^20, 28–29^. Particles were downsampled to a box size of 128 pixels (~2.67 Å/pixel) and used to train an eight-dimensional latent-variable model with 1024×3 encoder and decoder architectures. Particle poses and CTF parameters for CryoDRGN training were obtained from the non-uniform refinement. After 50 epochs of training, 100 volumes were sampled from the kmeans cluster centers of the latent embeddings. Visual inspection of the resulting volumes, guided by the corresponding atomic model, revealed structural variability in both the substrate density and its position relative to the ClpXP complex (Movie 1).

## Acknowledgments

This work was supported by startup funds from the Department of Biochemistry and Molecular Biophysics at Washington University School of Medicine, NIH grant R35-GM160328, and a Siteman Cancer Center Catalyst Award. Cryo-EM data were collected at the Washington University in St. Louis Center for Cellular Imaging (WUCCI). We thank members of the Ghanbarpour laboratory as well Tim Lohman and Eric Galburt for helpful discussions and feedback. We thank Joey Davis for valuable discussions during the early stages of this work. We thank SBGrid^44^ for providing the structural biology software packages used in this study.

## Author Contributions

Y.L. expressed and purified the proteins and performed all biochemical assays, except the ATPase assay and degradation assays conducted at different KCl concentrations, which were performed by N.I. A.G. and N.I. prepared samples for EM imaging and collected the data. A.G. processed the EM data, performed the reconstruction and refinement, built and refined the structural models, and wrote the first draft of the manuscript. All authors contributed to writing and editing the manuscript. A.G. supervised the project.

