## Supporting Information for "Electrostatic Complementarity at the ClpX Substrate-Entry Channel Governs ATP-Driven Protein Unfolding"

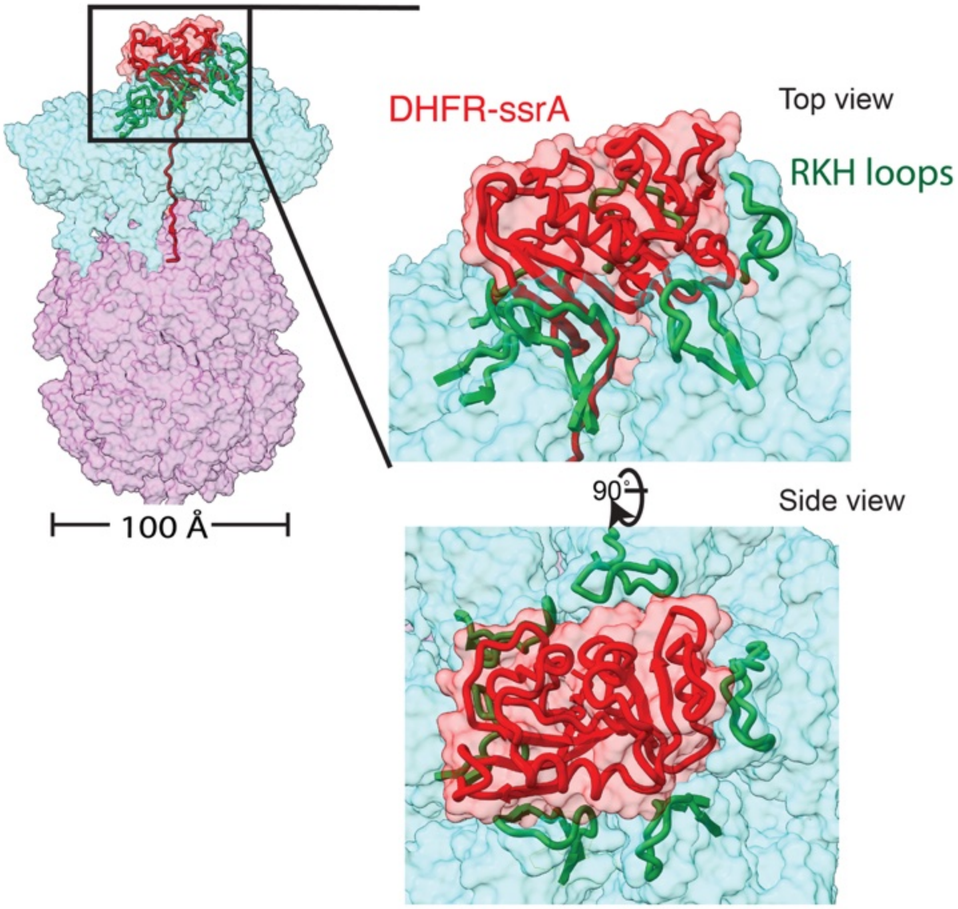


**Figure S1.** Cryo-EM structure of a substrate-engaged ClpX intermediate showing the folded DHFR–ssrA domain (red) positioned at the entrance of the axial channel and encircled by positively charged RKH loops (green) (PDB ID: 8V9R). ClpX is shown as a cyan surface and ClpP as a purple surface.

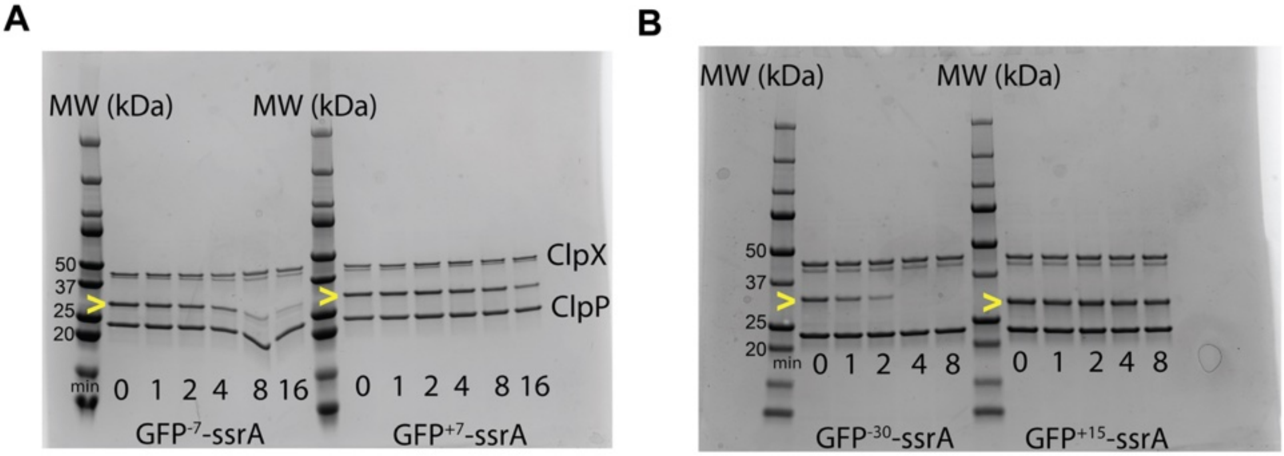


**Figure S2. SDS-PAGE gel degradation assays with GFP–ssrA variants.** SDS–PAGE analysis of ClpXP complexes incubated with different GFP–ssrA substrates over time. **(A)** $GFP^{-7}$–ssrA and $GFP^{+7}$–ssrA. **(B)** $GFP^{-30}$–ssrA and $GFP^{15+}$–ssrA. Samples were collected at the indicated time points and analyzed by SDS–PAGE. Bands corresponding to ClpX and ClpP are indicated. The yellow arrowheads mark substrate bands and their time-dependent disappearance during degradation.

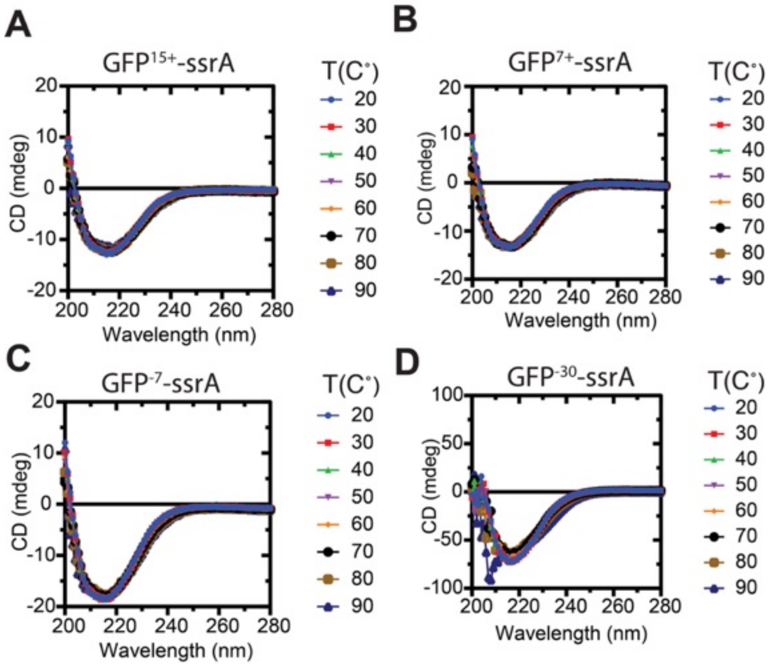


**Figure S3.** Temperature-dependent circular dichroism (CD) spectra of GFP–ssrA variants. CD spectra were recorded from 20–90 °C to evaluate the thermal stability and secondary-structure integrity of **(A)** GFP$^{15+}$–ssrA, **(B)** GFP$^{7+}$–ssrA, **(C)** GFP$^{-7}$–ssrA, and **(D)** GFP$^{-30}$–ssrA. Ellipticity changes in the far-UV region indicate temperature-induced structural perturbations, with negatively charged variants showing greater thermal sensitivity at elevated temperatures.

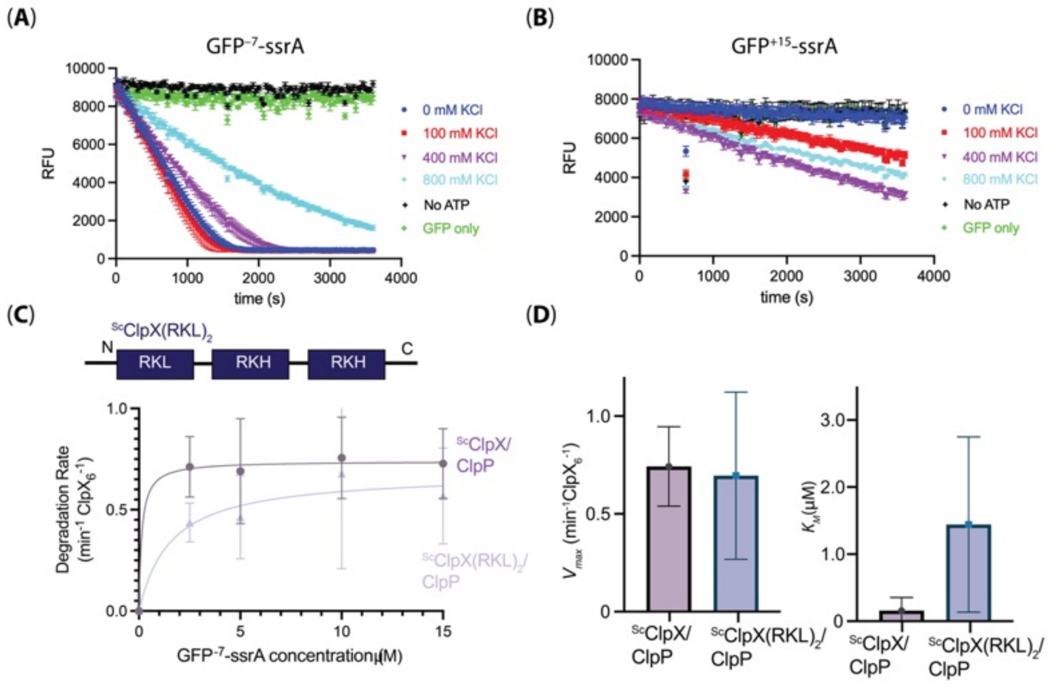


**Figure S4. Additional biochemical evidence supporting electrostatic contributions to ClpXP-mediated substrate degradation. (A,B)** Effect of ionic strength on degradation of charge-variant GFP substrates. **(A)** Increasing KCl concentration progressively reduced degradation of $GFP^{-7}$–ssrA. **(B)** Increasing KCl concentration from 0 to 400 mM enhanced degradation of $GFP^{+15}$–ssrA. No-ATP and GFP-only controls (100 mM KCl) are shown for comparison. Reactions contained 0.5 μM $ClpX_6^{\Delta N}$, 1.5 μM $ClpP_{14}$, and 10 μM GFP substrate. **(C,D)** Effect of RKL substitution in ClpX on degradation kinetics of $GFP^{-7}$–ssrA. **(C)** Schematic of the engineered ${}^{sc}ClpX^{\Delta N}(RKL)_2$ variant (top) and Michaelis–Menten analysis comparing degradation by ${}^{sc}ClpX^{\Delta N}$/ClpP and ${}^{sc}ClpX^{\Delta N}(RKL)_2$/ClpP over a range of $GFP^{-7}$–ssrA concentrations. **(D)** Corresponding kinetic parameters ($V_{max}$ and $K_M$) derived from the fits in **(C)**. Data represent the mean ± SD of three independent experiments.

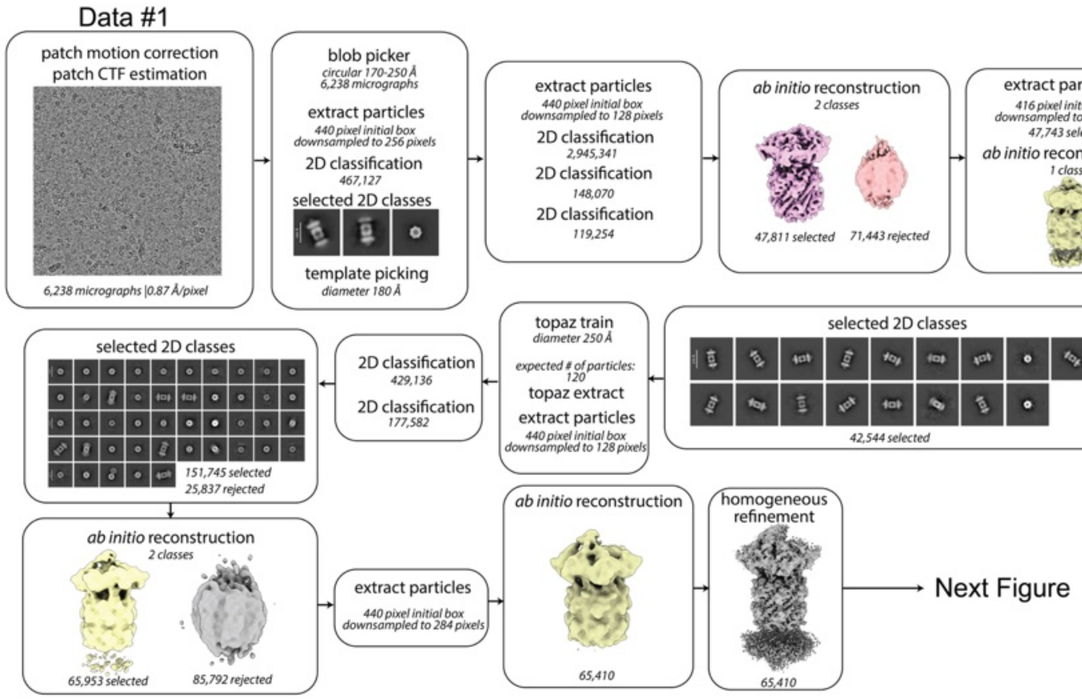
Data #1
patch motion correction
patch CTF estimation
6,238 micrographs |0.87 Å/pixel
blob picker
circular 170-250 Å
6,238 micrographs
extract particles
440 pixel initial box
downsampled to 256 pixels
2D classification
467,127
selected 2D classes
template picking
diameter 180 Å
extract particles
440 pixel initial box
downsampled to 128 pixels
2D classification
2,945,341
2D classification
148,070
2D classification
119,254
ab initio reconstruction
2 classes
47,811 selected
71,443 rejected
extract particles
416 pixel initial box
downsampled to 256 pixels
47,743 selected
ab initio reconstruction
1 class
homogeneous
refinement
47,743 selected
extract particles
640 pixel initial box
downsampled to 320 pixels
2D classification
47,704 selected
selected 2D classes
42,544 selected
topaz train
diameter 250 Å
expected # of particles:
120
topaz extract
extract particles
440 pixel initial box
downsampled to 128 pixels
2D classification
429,136
2D classification
177,582
selected 2D classes
151,745 selected
25,837 rejected
ab initio reconstruction
2 classes
65,953 selected
85,792 rejected
extract particles
440 pixel initial box
downsampled to 284 pixels
ab initio reconstruction
65,410
homogeneous
refinement
65,410
Next Figure

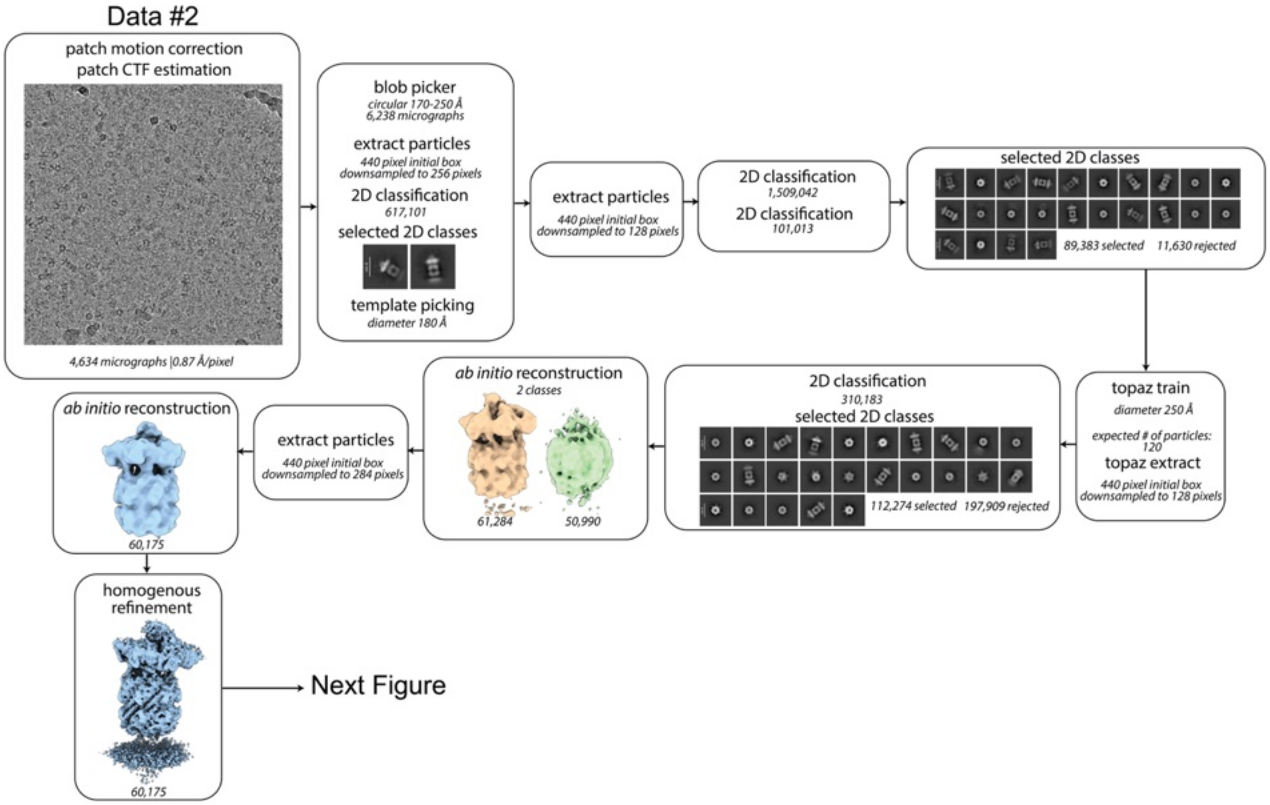
Data #2
patch motion correction
patch CTF estimation
4,634 micrographs |0.87 Å/pixel
blob picker
circular 170-250 Å
6,238 micrographs
extract particles
440 pixel initial box
downsampled to 256 pixels
2D classification
617,101
selected 2D classes
template picking
diameter 180 Å
extract particles
440 pixel initial box
downsampled to 128 pixels
2D classification
1,509,042
2D classification
101,013
selected 2D classes
89,383 selected
11,630 rejected
topaz train
diameter 250 Å
expected # of particles:
120
topaz extract
440 pixel initial box
downsampled to 128 pixels
2D classification
310,183
selected 2D classes
112,274 selected
197,909 rejected
ab initio reconstruction
2 classes
61,284
50,990
extract particles
440 pixel initial box
downsampled to 284 pixels
ab initio reconstruction
60,175
homogenous
refinement
60,175
Next Figure

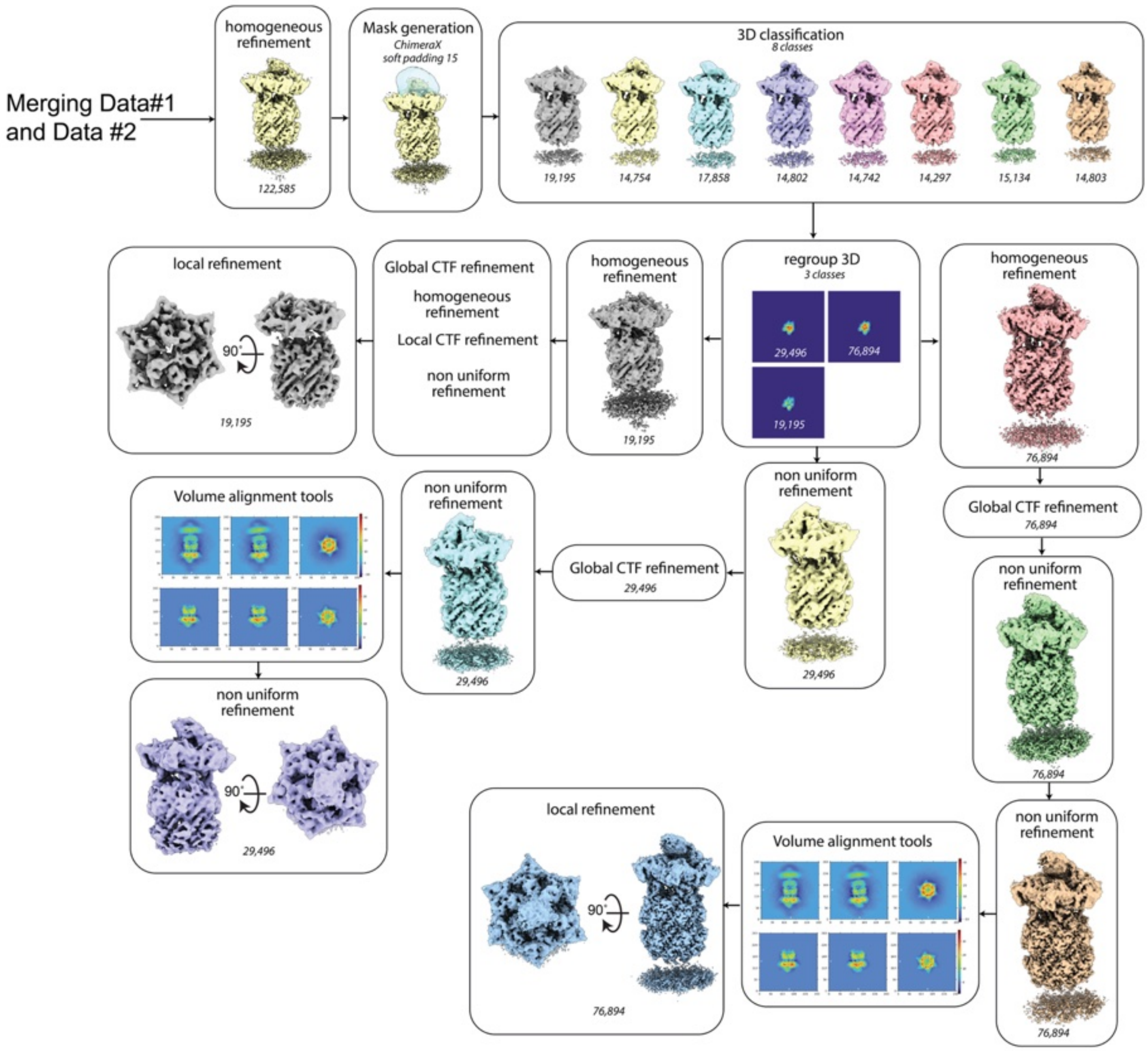


**Figure S5. CryoSPARC processing workflow for ClpX•ClpP•GFP$^{+15}$–ssrA complex.** Job names, job details, and non-default parameters (italicized) are noted in each box.

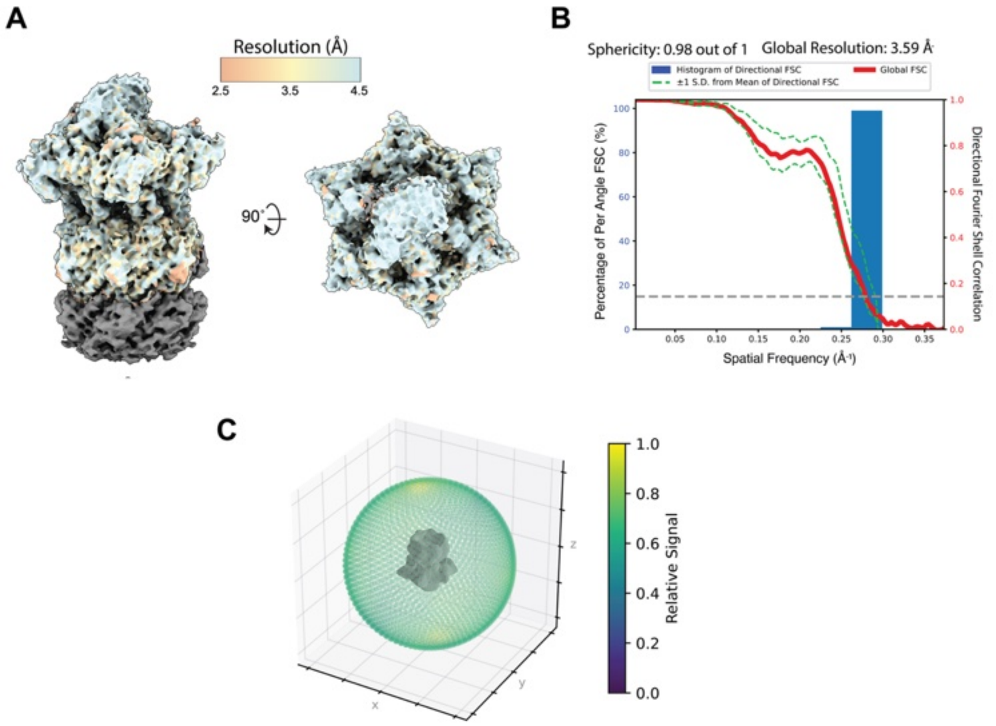


**Figure S6. Estimation of local and global resolution and angular distribution for ClpX•ClpP•GFP$^{+15}$–ssrA complex (intermediate I).** (**A**) Maps colored by local resolution as estimated by the cryoSPARC implementation of monoRes. (**B**) Global resolution and directional resolution calculated by 3DFSC server (https://3dfsc.salk.edu). (**C**) Projection angle distribution estimated by the cryoSPARC.

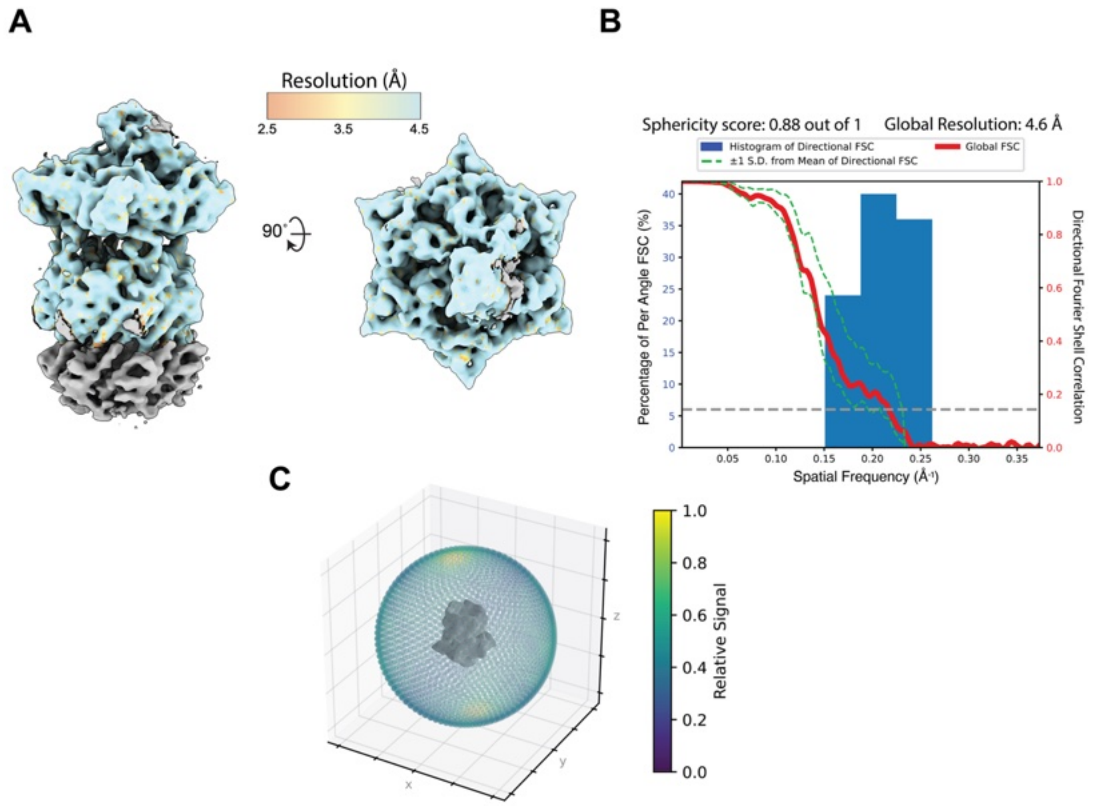


**Figure S7. Estimation of local and global resolution and angular distribution for ClpX•ClpP•GFP$^{+15}$–ssrA complex (intermediate II).** (**A**) Maps colored by local resolution as estimated by the cryoSPARC implementation of monoRes. (**B**) Global resolution and directional resolution calculated by 3DFSC server (https://3dfsc.salk.edu). (**C**) Projection angle distribution estimated by the cryoSPARC.

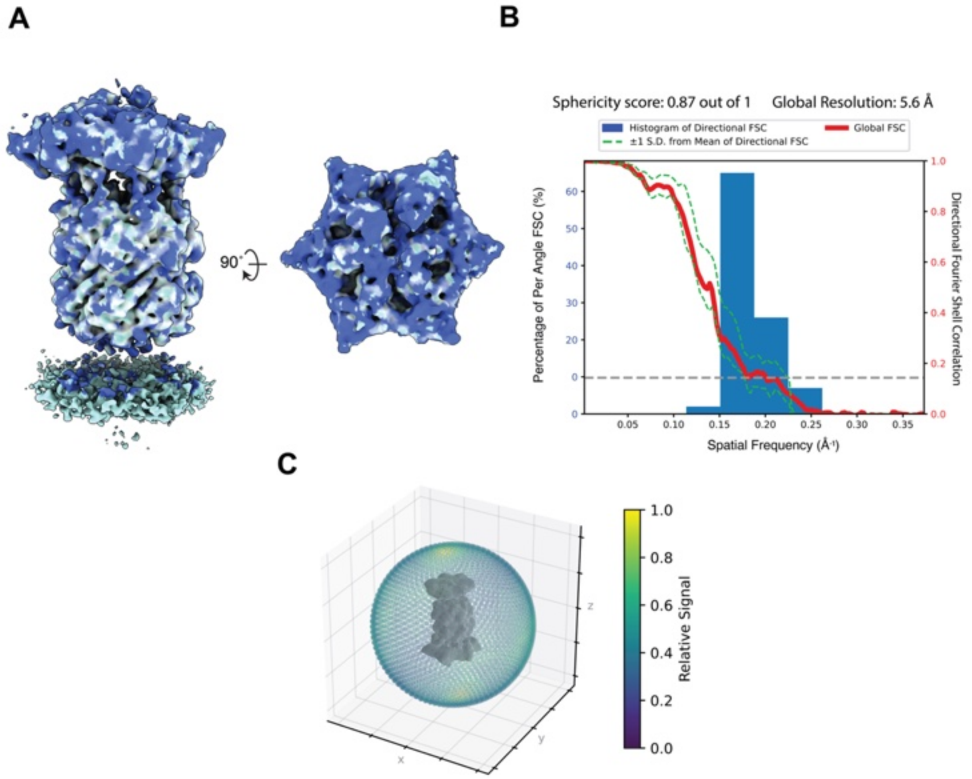


**Figure S8. Estimation of local and global resolution and angular distribution for ClpX•ClpP•GFP$^{+15}$–ssrA complex (intermediate III).** (**A**) Maps colored by local resolution as estimated by the cryoSPARC implementation of monoRes. (**B**) Global resolution and directional resolution calculated by 3DFSC server (https://3dfsc.salk.edu). (**C**) Projection angle distribution estimated by the cryoSPARC.

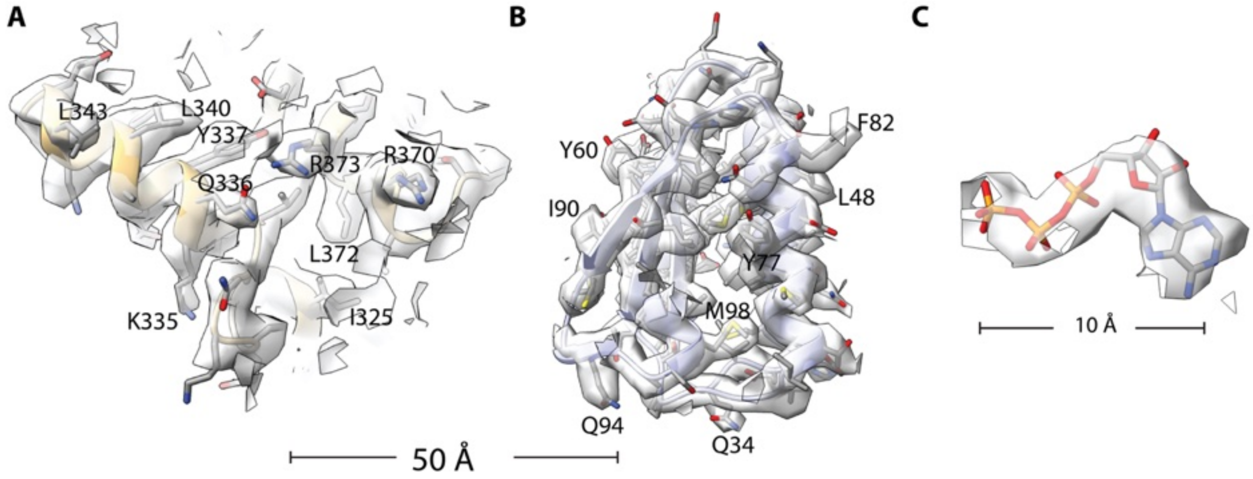


**Figure S9. Representative cryo-EM density overlays for intermediate I of the ClpX•ClpP•GFP$^{+15}$–ssrA complex.** (**A**) Local density for ClpX chain A (residues 320–380). (**B**) Density corresponding to ClpP chain H (residues 1–100). (**C**) Density for ATP bound to ClpX chain A. Scale bars, 50 Å (**A,B**) and 10 Å (**C**). The density map was sharpened using DeepEMhancer in CryoSPARC.

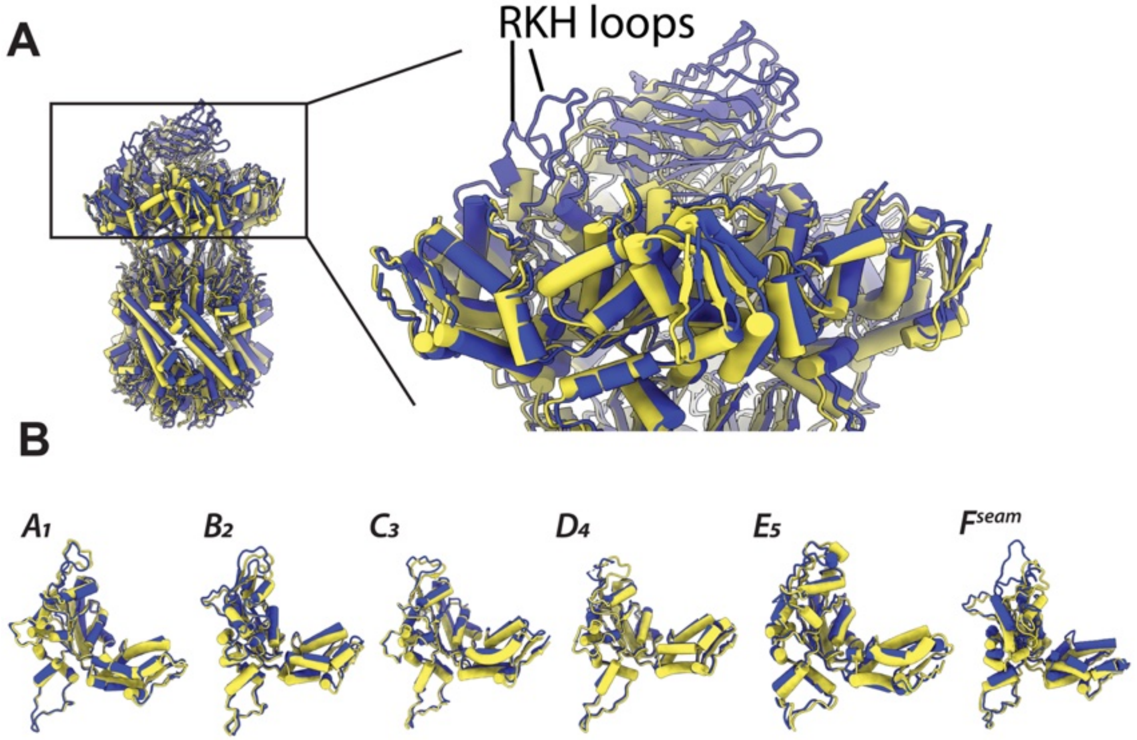


**Figure S10. Structural comparison of substrate-bound ClpXP complexes.** (**A**) Overlay of cryo-EM structures of ClpXP engaged with different substrates: GFP$^{+15}$–ssrA (blue) and DHFR–ssrA bound to Methotrexate (PDB: 8V9R; yellow), highlighting overall conformational similarity of ClpX. The boxed region is enlarged to emphasize differences in substrate engagement and RKH loop interactions. Structures are shown in cartoon representation with the two complexes colored in blue and yellow. (**B**) Superposition of individual ClpX subunits ($A_1$–$E_5$ and seam subunit) from the aligned complexes.

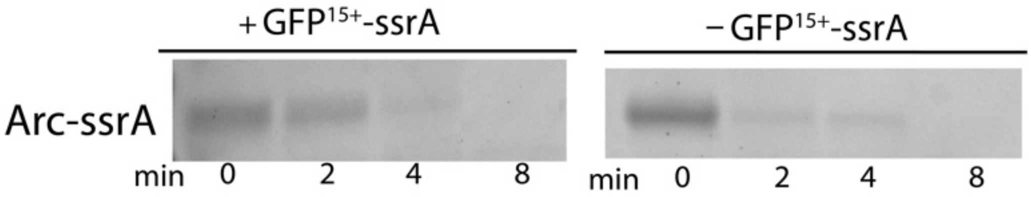


**Figure S11. Effect of ClpXP pre-engagement with GFP$^{+15}$–ssrA on subsequent substrate degradation.** ClpXP complexes were pre-incubated with (+GFP$^{+15}$–ssrA) or without (−GFP$^{+15}$–ssrA) GFP$^{+15}$–ssrA in the presence of 5 mM ATP at 30 °C and then diluted 10-fold into reactions containing 10 μM Arc–ssrA (total volume: 40μL). Arc–ssrA degradation was monitored by SDS-PAGE at 0, 2, 4, and 8 min. Pre-engagement with GFP$^{+15}$–ssrA slowed the initial rate of Arc–ssrA degradation but did not prevent complete substrate turnover, indicating that GFP$^{+15}$–ssrA does not stably trap or irreversibly stall the ClpXP protease. The 10X pre-engagement reaction contained 5 μM ClpX, 5 μM ClpP, 5 μM GFP$^{+15}$–ssrA, and 5 mM ATP, whereas the control reaction contained 5 μM ClpX, 5 μM ClpP, and 5 mM ATP.

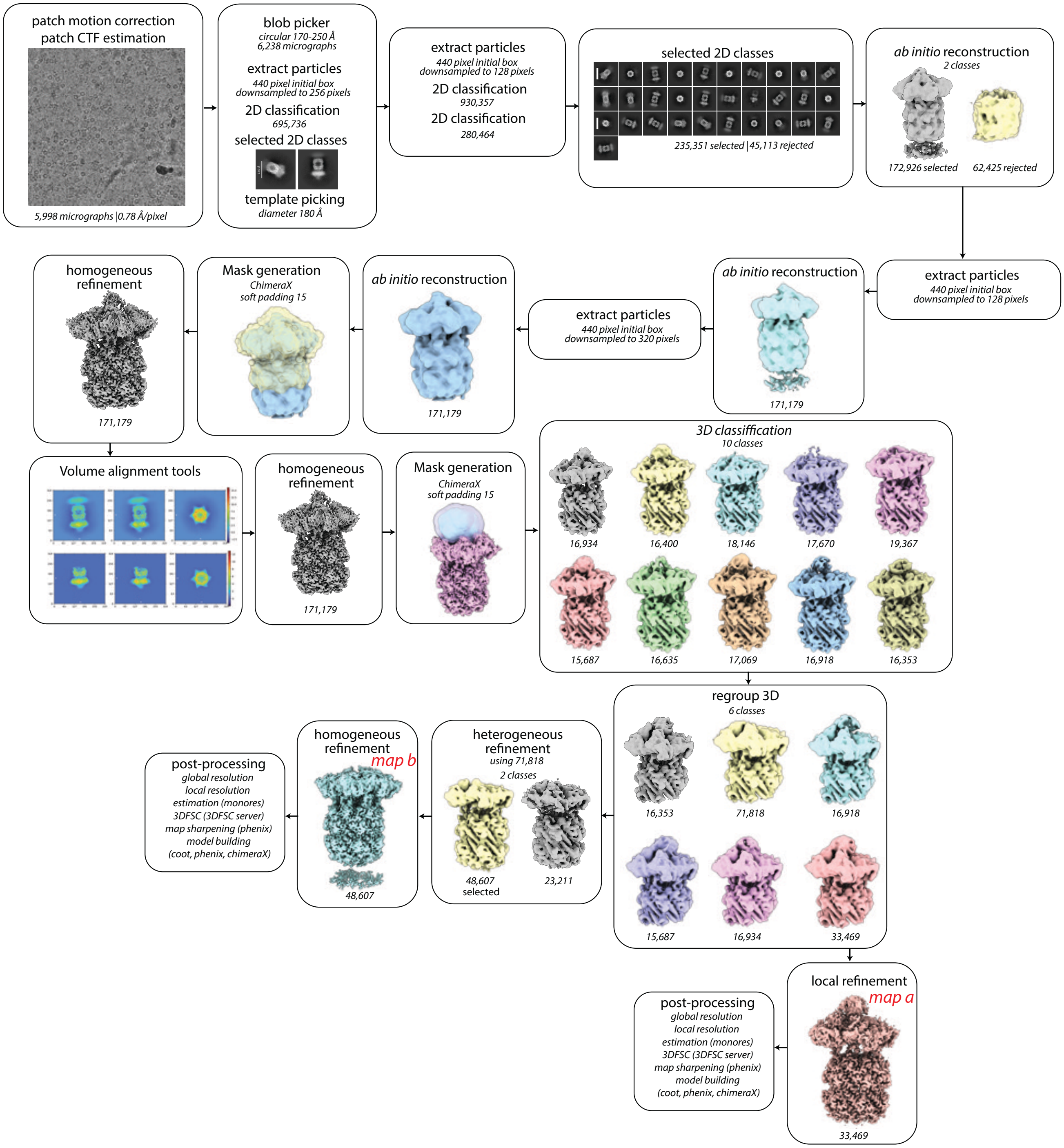


**Figure S12. CryoSPARC processing workflow for ClpX•ClpP•GFP$^{-7}$–ssrA.** Job names, job details, and non-default parameters (italicized) are noted in each box.

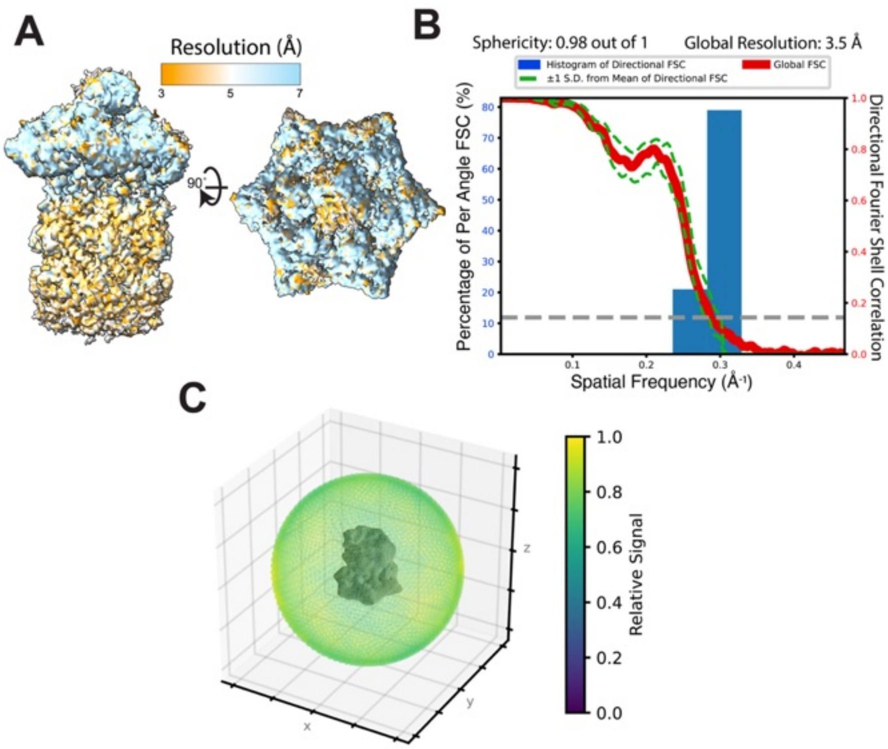


**Figure S13. Estimation of local and global resolution and angular distribution for ClpX•ClpP•GFP-7–ssrA (map a).** (**A**) Maps colored by local resolution as estimated by the cryoSPARC implementation of monoRes. (**B**) Global resolution and directional resolution calculated by 3DFSC server (https://3dfsc.salk.edu). (**C**) Projection angle distribution estimated by the cryoSPARC.

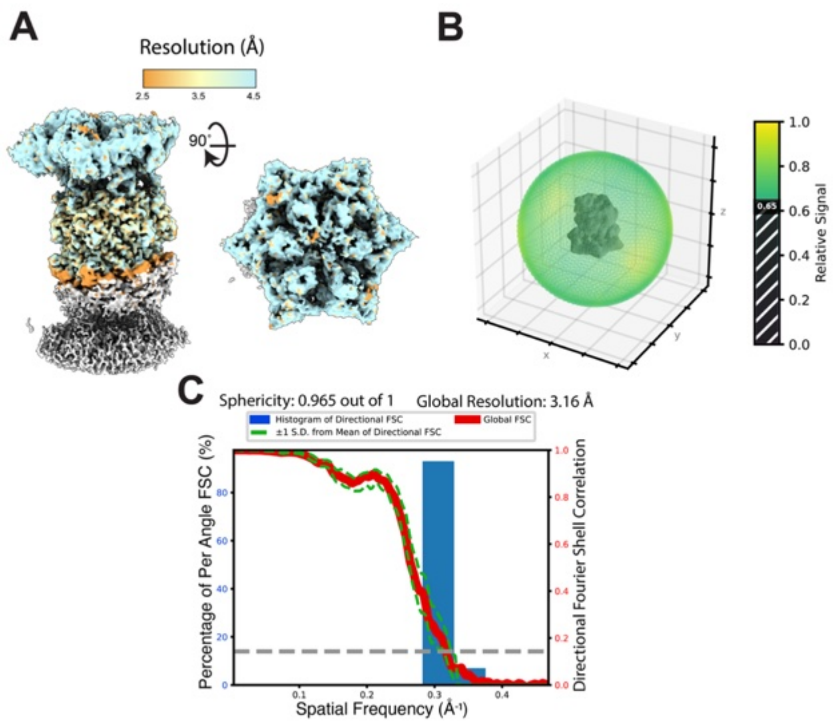


**Figure S14. Estimation of local and global resolution and angular distribution for ClpX•ClpP•GFP-$^{7}$–ssrA (map b).** (**A**) Maps colored by local resolution as estimated by the cryoSPARC implementation of monoRes. (**B**) Projection angle distribution estimated by the cryoSPARC. (**C**) Global resolution and directional resolution calculated by 3DFSC server (https://3dfsc.salk.edu).

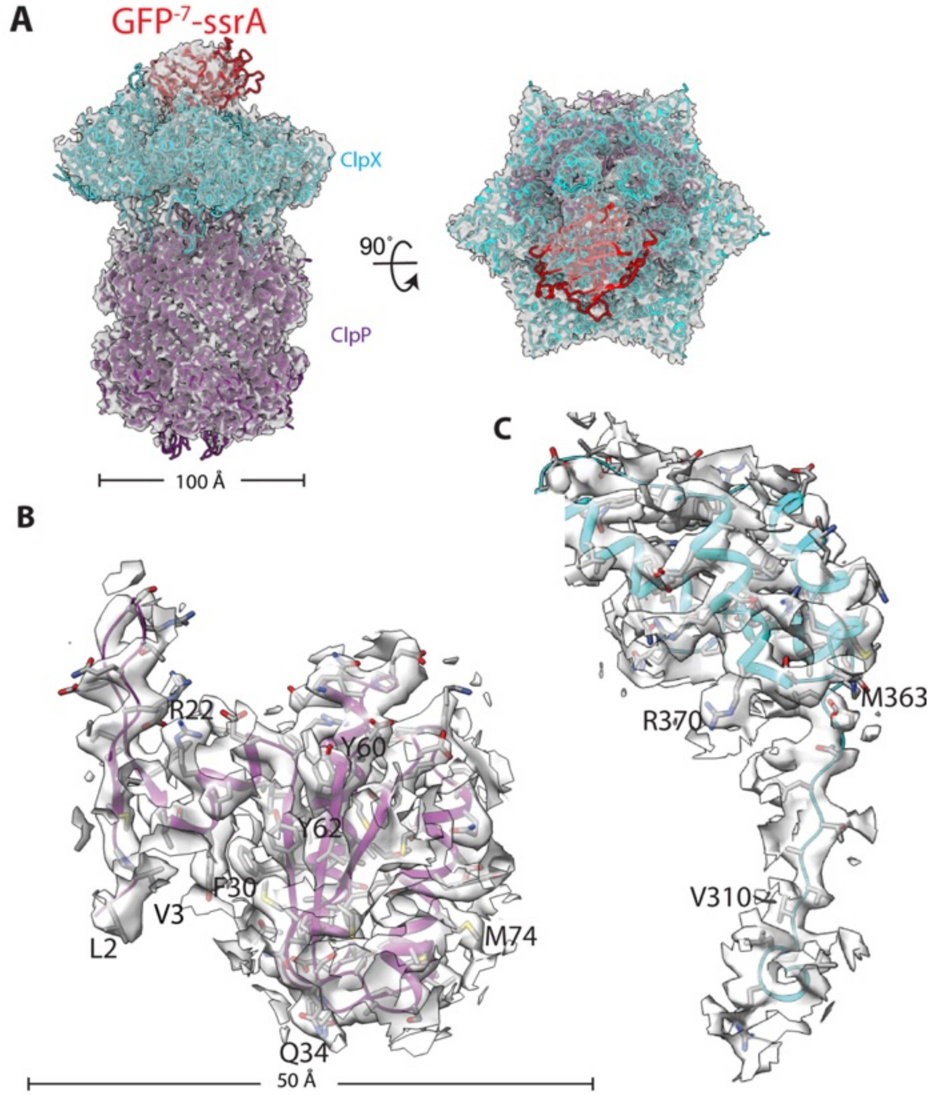


**Figure S15. Cryo-EM reconstruction of the GFP$^{-7}$–ssrA•ClpXP complex. The density map was sharpened using CryoSPARC.** (**A**) Overall cryo-EM density map of the GFP$^{-7}$ssrA•ClpXP complex with fitted atomic models shown in ribbon representation. GFP$^{-}$7–ssrA is colored red, ClpX cyan, and ClpP purple. Two orthogonal views are presented. Scale bar, 100 Å. (**B**) Local density for ClpP chain H (residues 1–100). (**C**) Local density for ClpX chain A (residues 305–380). Scale bars, 50 Å.

**Table S1**. Cryo-EM data collection, processing, model building, and validation statistics.

| Name | ClpXP-GFP$^{+15}$–ssrA (intermediate I) | ClpXP-GFP$^{+15}$–ssrA (intermediate II) | ClpXP-GFP$^{+15}$–ssrA (intermediate III) | ClpXP-GFP$^{-7}$–ssrA (map a) | ClpXP-GFP$^{-7}$–ssrA (map b) |
|---|---|---|---|---|---|
| PDB ID | 37DM | NA | NA | 36ON | 37DI |
| EMDB ID | EMD-78093 | EMD-78092 | EMD-78091 | EMD-77726 | EMD-78089 |
| Microscope | Titan Krios G3 | Titan Krios G3 | Titan Krios G3 | Titan Krios G3 | Titan Krios G3 |
| Camera | Falcon 4 | Falcon 4 | Falcon 4 | Falcon 4 | Falcon 4 |
| Magnification | 75,000 | 75,000 | 75,000 | 130,000 | 130,000 |
| Voltage (kV) | 300 | 300 | 300 | 300 | 300 |
| Total electron dose (e-/Å2) | 51.8 $e^-$/Å2 | 51.8 $e^-$/Å2 | 51.8 $e^-$/Å2 | 55.48 $e^-$/Å2 | 55.48 $e^-$/Å2 |
| Defocus range (µm) | −0.4 to −2.0 µm | −0.4 to −2.0 µm | −0.4 to −2.0 µm | −0.4 to −2.0 µm | −0.4 to −2.0µm |
| Pixel size (Å) | 0.87 | 0.87 | 0.87 | 0.78 | 0.78 |
| Micrographs collected | 4634 | 4634 | 4634 | 6000 | 6000 |
| Final particles | 76894 | 29,496 | 29,496 | 33,469 | 48,607 |
| Symmetry | C1 | C1 | C1 | C1 | C1 |
| Resolution (Å) (0.143) | 3.35 | 4.48 | 4.48 | 3.14 | 2.90 |
| Unmasked resolution (Å) | 5.7 | 7.8 | 7.8 | 5.8 | 4.1 |
| Global resolution (Å) | 3.59 | 4.64 | 5.60 | 3.48 | 3.16 |
| All atoms | 78468 | NA | NA | 77802 | 73613 |
| Protein residues | 5007 | NA | NA | 4973 | 4723 |
| Ligands | ATP:3; $Mg^{+2}$:4; ADP:3 | NA | NA | ATP:3 $Mg^{+2}$:4; ADP: 3 | ATP:3; $Mg^{+2}$:4; ADP:3 |
| Map-model CC (mask/box/peaks) | 0.75/0.82/0.70 | NA | NA | 0.64/0.69/0.56 | 0.73/0.72/0.62 |
| RMSD bond lengths (Å) | 0.004 | NA | NA | 0.003 | 0.004 |
| RMSD bond angles (deg.) | 0.722 | NA | NA | 0.584 | 0.722 |
| MolProbity score | 1.1 | NA | NA | 1.32 | 0.97 |
| Clash score | 3.35 | NA | NA | 5.85 | 2.04 |
| C-beta outliers (%) | NA | NA | NA | NA | NA |
| Rotamer outliers (%) | 0.9 | NA | NA | 0.6 | 0.9 |
| Ramachandran favored (%) | 99.82 | NA | NA | 99.96 | 99.90 |
| Expected Q-score | 0.48 | NA | NA | 0.49 | 0.55 |
| ClpX Q-score | 0.50 | NA | NA | 0.42 | 0.58 |
| ClpP Q-score | 0.50 | NA | NA | 0.42 | 0.58 |
| Substrate Q-score | 0.50 | NA | NA | 0.42 | 0.58 |
